# Maternal care and early rearing environment influence puppy behaviour and cognition

**DOI:** 10.64898/2025.11.29.691282

**Authors:** Emily E. Bray, Laura E.L.C. Douglas, Kerinne M. Levy, Gitanjali E. Gnanadesikan, Daniel J. Horschler, Brenda S. Kennedy, Evan L. MacLean

## Abstract

Relatively little is known about the influence of maternal care on dog behaviour, despite the prominence of dogs in our lives and the established importance of early experiences in other species. While recent observational research has begun to document associations between canine maternal behaviour and later offspring outcomes, there is still much to learn—particularly regarding what factors impact maternal behaviour and the enduring effects of maternal care on puppy behavioural development. Understanding how early experiences shape future behaviour is of practical importance for companion and working dogs. We characterized the early rearing environment of 235 individual puppies from 59 litters bred by a service dog provider and explored whether offspring cognitive and behavioural traits through 16 months of age were associated with early mothering behaviour. We also investigated whether maternal behaviour could be predicted by pre-pregnancy dam behaviour and how the rearing location (private home or professional breeding centre) influenced maternal behaviour and/or puppy behaviour. We identified dam behavioural characteristics measured pre-pregnancy that were related to subsequent maternal care. While time of year was associated with maternal behaviour scores, parity, litter size, breed composition, and rearing location were not. We found that rearing location was related to puppy performance on the Dog Cognitive Development Battery (DCDB) at 8 weeks of age. Finally, maternal behaviour was associated with certain puppy cognitive and behavioural measures, assessed via the DCDB at 8 weeks of age and via questionnaires at 6, 10, 12 and 16 months of age. Our results indicate that experiences within the first few weeks of life, particularly maternal care and rearing location, may influence several aspects of dog behaviour relevant to both working and companion animals.

**Highlights:**

- We tracked early maternal interactions across 59 litters
- Some traits measured pre-pregnancy were associated with subsequent maternal care
- Rearing location was associated with some puppy behaviour but not maternal care
- Maternal care was associated with some aspects of puppy behaviour at 8 weeks
- Maternal care was associated with reports of puppy behaviour through 16 months

## Introduction

Early life experiences are crucial across many animal species, shaping development by programming biological systems and laying the foundation for long-term impacts. In mammals particularly, the mother plays a prominent role in these early life interactions, with cascading effects. The rodent and primate literature are replete with examples of how early maternal care impacts offspring far into the future (Maestripieri, 2018). For example, in rodent models, early-life stressors ranging from maternal separation to altered maternal behaviour caused by limited nesting material have been shown to induce lasting changes in progeny behaviour, stress reactivity, and neurodevelopment (Kaffman & Meaney, 2007; Walker et al., 2017).

However, in comparison and despite their prominence in our lives, we know relatively little about the impact of mothering behaviours on offspring behaviour in dogs (Li, 2024). Understanding the early environmental factors that influence dog behaviour later in life is of practical importance for both working dogs and companion dogs. For example, identifying maternal behaviours that contribute to desirable behaviour in offspring could inform breeding plans, thereby conserving resources and serving more clients at working dog organizations (Bray, Otto, et al., 2021; Foyer et al., 2016; Li, 2024). Optimizing the early environment to support desirable companion dog behaviours could potentially result in fewer pet dogs being relinquished for behavioural problems (Czerwinski et al., 2016; Li, 2024; Santos, Beck, & Fontbonne, 2020).

Within the past decade, observational studies have begun to describe and characterize variation in early maternal behaviour in dogs and then explore association with later offspring phenotypes (Baqueiro-Espinosa et al., 2022; Baqueiro-Espinosa et al., 2025; Bray et al., 2017b; Foyer et al., 2016; Guardini et al., 2017; Guardini et al., 2016; Montgomery et al., 2025). These studies focus on maternal care during the neonatal and transition periods—the first 21 days following birth—when puppies are still extremely dependent on their mother and maternal care may be particularly influential to their subsequent development (Santos, Beck, & Fontbonne, 2020).

Maternal care has most often been characterized by behavioural measures of proximity (amount of time spent near or in contact with puppies), oronasal interactions (time spent licking, sniffing, or grooming), and feeding (time spent nursing) (Baqueiro-Espinosa et al., 2024; Baqueiro-Espinosa et al., 2022; Bray et al., 2017a; Foyer et al., 2016; Guardini et al., 2017; Guardini et al., 2015; Guardini et al., 2016; Santos, Beck, Blondel, et al., 2020). Our previous research (Bray et al., 2017a, 2017b) included vigilance behaviours (orienting out of the whelping area) as a component of maternal care as well.

Thus far, research in this area has revealed some consistent patterns related to maternal care. For example, the overall level of maternal behaviour decreases over time during the first three weeks as puppies gain independence (Baqueiro-Espinosa et al., 2024; Bray et al., 2017a; Foyer et al., 2016; Guardini et al., 2017; Guardini et al., 2015; Guardini et al., 2016; Montgomery et al., 2025; Santos, Beck, Blondel, et al., 2020). The only tracked behaviour that does not decrease over time is vertical nursing, which starts out relatively rare and increases as the puppies get older (Baqueiro-Espinosa et al., 2024; Bray et al., 2017a; Montgomery et al., 2025). Despite a decrease in the amount of overall maternal care as puppies age, a dam’s maternal behaviour remains relatively stable across weeks, and there are distinct variations in the style and quantity of maternal care between dams (Bray et al., 2017a; Foyer et al., 2016;

Montgomery et al., 2025). Even in free-ranging dogs, where young pups are solely dependent on intraspecific care with no human caregiver support, researchers have identified similar patterns. For example, free-ranging dog mothers show individual differences in their maternal behaviour (Pal et al., 2021; Pal, 2005), and the quantity of their direct care (e.g., nursing, grooming, physical contact) declines over time as the puppies age (Pal et al., 2021; Pal, 2005; Paul & Bhadra, 2018; Paul et al., 2017).

Variation in maternal care might be partially explained by demographic factors, including breed, litter size, and parity. For example, our research found that Labrador retrievers had significantly higher maternal behaviour scores compared to German shepherds, indicating they were more often present and interactive with their puppies, and specifically displayed more contact per puppy (Bray et al., 2017a). Baqueiro-Espinosa et al. (2022) compared maternal behaviour by breed size, finding that small-breed dams spent more time in contact with and nursing puppies than medium- and large-breed dogs. Differences in care by breed size may be partially mediated by litter size, as smaller breed dogs tend to have smaller litters (Borge et al., 2011), and there is evidence that litter size might influence maternal care (Baqueiro-Espinosa et al., 2024; Bray et al., 2017a; Foyer et al., 2016; Foyer et al., 2013; Paul et al., 2017). Existing research with large breed dogs has also found that dams with smaller litters have higher maternal behaviour scores and spend more time in contact per puppy (Bray et al., 2017a), spend more time in the whelping box (Baqueiro-Espinosa et al., 2024) and have higher mother-puppy interaction scores (Foyer et al., 2016). Although in contrast, Baqueiro-Espinosa et al. (2024) found dams spent more total time nursing as the litter size increased, and Montgomery et al. (2025) found no effect of litter size on maternal behaviour. Finally, the effect of parity on maternal care has been inconsistent and needs further exploration (Santos, Beck, & Fontbonne, 2020). For example, Guardini et al. (2015) and Bray et al. (2017a) reported that less experienced mothers exhibited higher levels of maternal care, while Foyer et al. (2016) found no effect of parity on maternal care, and Montgomery et al. (2025) reported an interaction between parity and delivery method on maternal care (2024).

While maternal care may be affected by breed, litter, and parity, we have limited control over these variables, and they do not account for all the behavioural variation amongst dams. Thus, one worthwhile question is whether there are individual factors that we can reliably measure to predict what kind of mother a dam will be. Only two studies to date have explored whether a dam’s maternal behaviour can be predicted by her behaviour pre-pregnancy (Baqueiro-Espinosa et al., 2024; Montgomery et al., 2025). Baqueiro-Espinosa et al. (2024) evaluated dams on a behaviour assessment during the third week of gestation and before an enrichment protocol began. The only maternal behaviour that was related to the prenatal behaviour assessment was average time spent licking puppies. Specifically, dams who were scored on a behavioural test as more sociable, less fearful and reactive, and more willing to interact with novel stimuli spent less time licking their puppies. Using C-BARQ scores, Montgomery et al. (2025) found that dams who displayed more human-oriented separation-related behaviours exhibited lower levels of maternal care, and dams rated as more excitable spent more time in a vertical nursing position. Another set of environmental factors that might impact maternal behaviour are attributes of the physical location where the dam whelps and rears her offspring, but to our knowledge no studies have directly observed and compared the maternal behaviour of dams across different whelping environments. For working dog organizations and companion dog breeders, examining these attributes may provide additional insight into where and how to invest resources toward the physical spaces where dams whelp and rear their litters.

What about the impact of aspects of the early environment on puppy behaviour? Research comparing the effects of the early rearing environment on puppy behaviour exists, but is sparse, and has found some evidence for behavioural differences between dogs reared in a domestic home environment and those reared in a kennel environment during the first weeks of life (Appleby et al., 2002; Lenkei et al., 2019; Majecka et al., 2020; Wright, 1983).

There has been only slightly more research regarding how maternal behaviour impacts the behaviour of progeny as they develop. In particular, three studies have looked at the impact of early maternal care on puppy behaviour at 8 weeks of age: two studies found higher maternal care was linked to more confident (Guardini et al., 2016) and sociable (Baqueiro-Espinosa et al., 2025) behaviour in puppies reared in a commercial breeding kennel, while a third study found differing results in that high maternal care was linked to less willingness to engage along with more stress behaviours in puppies reared in private homes (Guardini et al., 2017). Montgomery et al. (2025) assessed puppies in a detection dog program at 12 weeks of age and found no effect of maternal care on scores on a behavioural test assessing reward motivation, detection ability, and environmental soundness. The differences between these studies raise questions about how the relationship between maternal care and offspring behaviour potentially differs between populations of dogs, as well as how it combines with other aspects of the early environment to affect later puppy characteristics.

Additional research suggests maternal care may have even longer-lasting effects on puppies’ behaviour and temperament. In a questionnaire-based study of dog owners, fearful and anxious behaviours in adult dogs—including noise sensitivity and separation anxiety—were associated with reported poor maternal care and being reared by a dam who spent less time with the puppies (Tiira & Lohi, 2015). In a study of military working dogs, higher levels of maternal care were associated with higher levels of behaviours categorized as Social Engagement, Physical Engagement, and Aggression, measured when dogs were 15-18 months old (Foyer et al., 2016). Finally, in our own research with guide dogs, we similarly found maternal care was related to the behaviour of offspring evaluated after one year of age, and was associated with some undesirable behaviours, and ultimately failure to succeed as a guide dog (Bray et al., 2017b). Specifically, dogs who experienced high levels of maternal care demonstrated higher activity levels while in isolation, a shorter latency to vocalize when introduced to a novel object, and less competence during problem-solving tasks. Similarly, recent research (Montgomery et al., 2025) found dogs who experienced more maternal care were also less likely to be selected as detection dogs.

Thus, while the existing research suggests maternal care has a long-term influence on puppy development, more research is needed to clarify the specific impact. Effects may vary by breed, rearing environment (in-home or professional facility), and puppy age at evaluation, among other variables. The existing research on maternal care and working dogs suggests that a higher level of maternal care is not always better, and that the optimal amount of maternal care may be relative to the population (Bray et al., 2017b; Montgomery et al., 2025). Thus, studying maternal care in additional populations of working dogs will help to elucidate which levels are optimal for different working dog roles.

By characterizing canine maternal care and measuring subsequent puppy outcomes, here we aim to address several gaps and supplement the existing literature on associations between maternal care and puppy development. Most previously published observations of maternal behaviour have been conducted in a professional kennel environment, either at a working dog (Bray et al., 2017a, 2017b; Foyer et al., 2016; Montgomery et al., 2025; Santos, Beck, Blondel, et al., 2020) or commercial breeding (Baqueiro-Espinosa et al., 2024; Baqueiro-Espinosa et al., 2022; Guardini et al., 2016) facility, and no research to our knowledge has directly compared maternal behaviour between professional facilities and private home rearing environments within a population of dogs. Studies have typically enrolled 30 dams or fewer (with the exception of Santos, Beck, Blondel, et al., 2020), and several studies recorded maternal behaviour during the morning only (Guardini et al., 2017; Guardini et al., 2015; Guardini et al., 2016). Importantly, with the exception of Guardini et al. (2017; 2015; 2016), all previous studies have been unable to distinguish between individual puppies, instead assigning all littermates the same maternal care score. Finally, puppy behaviour has typically been evaluated using only a single type of evaluation in any given study: either a questionnaire (Tiira & Lohi, 2015) or a behaviour test, such as an arena or isolation test (Guardini et al., 2017; Guardini et al., 2015; Guardini et al., 2016) or a battery of cognitive and/or temperament tasks (Bray et al., 2017a, 2017b; Foyer et al., 2016; Montgomery et al., 2025).

In the current study, we enrolled 235 individual puppies from 59 litters, split evenly between those who were whelped and reared in a private home and those who were whelped and reared in a professional centre. We observed the dogs over the first three weeks post-whelp and characterized the maternal behaviour provided by dams. We then explored if cognitive and behavioural traits were associated with early environmental features using both behavioural questionnaires and a test battery with cognitive and temperament tasks. Specifically, we asked if pre-pregnancy dam behavioural characteristics predicted the type of maternal behaviour she would ultimately display, if rearing location was related to maternal behaviour and/or puppy behaviour, and if early maternal behaviour was associated with later puppy behaviour and cognition.

## General Methods

### Ethical Note

All procedures were approved and adhered to regulations set forth by the Institutional Animal Care and Use Committee at the University of Arizona (IACUC No. 16-175). The behavioural tasks in which dams and puppies participated were designed to be non-invasive. Dogs ate their usual kibble over the course of the testing session, ensuring all participants were fed their usual amount. We incorporated plenty of play and bathroom breaks to ensure the testing experience was a positive one, and every task involved food rewards, play, and/or praise. We also adhered to strict abort criteria that allowed dogs to opt out of a task if they were no longer engaged.

### Timeline and Study Design

All data was collected from dogs at Canine Companions, the largest service dog provider in the United States. Our overall study design is described in Figure 1. Prospective dams first participated in cognitive and temperament testing—the Dog Cognitive Development Battery (DCDB; Bray et al., 2020)—when they were in oestrus, prior to becoming pregnant with the study litter (with the exception of six dams, who were tested after weaning). This testing occurred between November 2017 and May 2022. In-season females are boarded at Canine Companions headquarters for the duration of their oestrus cycle, which allowed us to collect these measures. We also opportunistically collected behavioural information reported by these dogs’ puppy raisers when they were approximately 1-year old, prior to returning for professional training and ultimately being selected as breeders.

**Figure 1.**
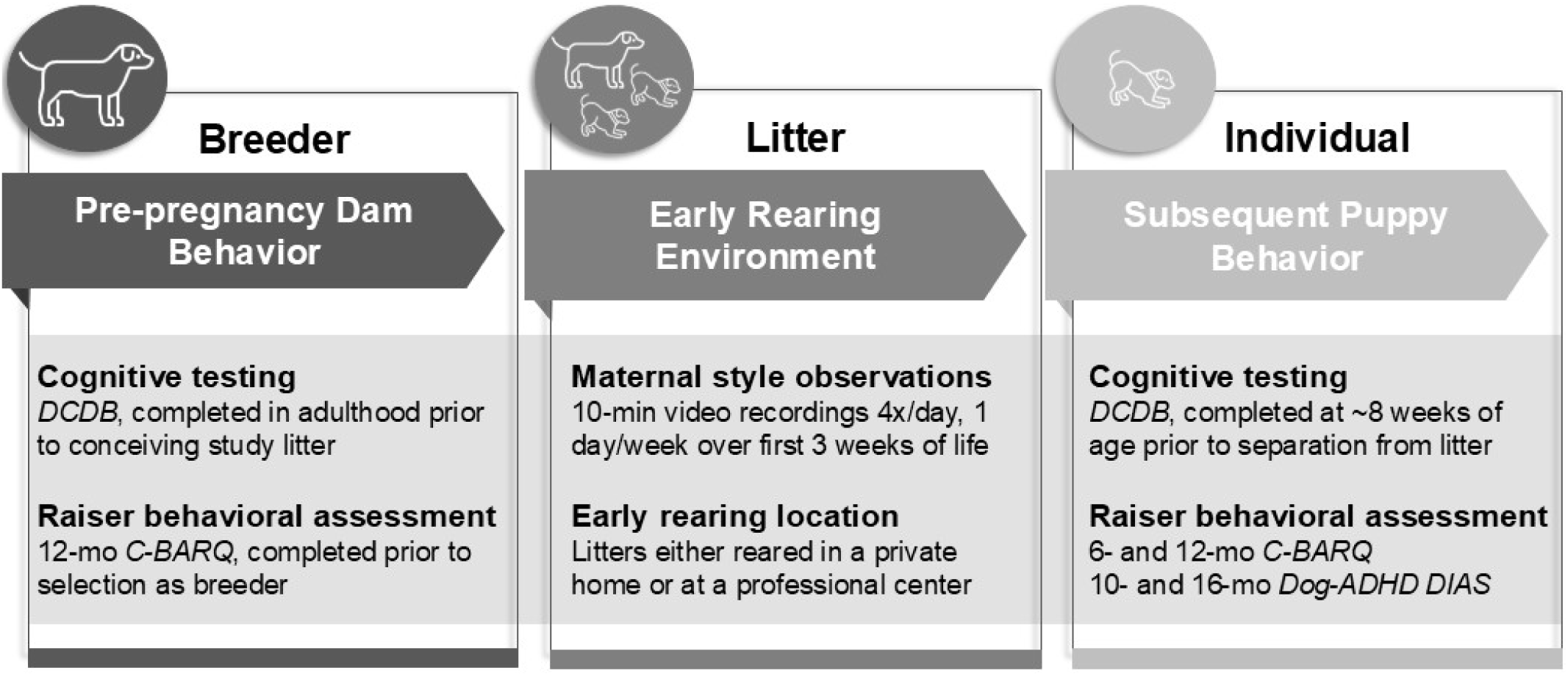
Data collection study design. DCDB = Dog Cognitive Development Battery, C-BARQ = Canine Behavioral Assessment & Research Questionnaire, Dog-ADHD = adapted human attention deficit hyperactivity disorder questionnaire, DIAS = Dog Impulsivity Assessment Scale.

After the two-month gestation period, dams whelped and video observations began. We recruited dams who would be rearing their puppies in one of two different environments: half of the dams were with their puppies in a volunteer breeder caretaker home (“BC dams”), while the other half of the dams were with their puppies at the Canine Early Development Center, a professional facility at Canine Companions (“CEDC dams”). Our goal was to enrol dams equally across the two different locations to allow for comparison of early environments. To avoid introducing biases, we enrolled BC and CEDC litters at roughly the same rate. The first study litter whelped in January 2018 and litters were enrolled through February 2021. We enrolled 31 litters (126 puppies) whelped in 2018, 14 litters (54 puppies) whelped in 2019, 13 litters (51 puppies) whelped in 2020, and 1 litter (4 puppies) whelped in early 2021. When considering all years of the study, we enrolled at least 12 litters in each season. Specifically, we enrolled 12 litters (50 puppies) whelped in January – March, 19 litters (73 puppies) whelped in April – June, 12 litters (48 puppies) whelped in July – September; and 16 litters (64 puppies) whelped in October – December. The entire enrolment timeline is specified in the supplemental materials (Data_collection_timeline tab).

Each litter was video recorded and the maternal behaviour of the dam toward her litter was observed over the first three weeks post-whelp while the litter was still contained in the whelping pool or box. We also collected biological samples from the puppies at two separate timepoints between 6 and 7.5 weeks of age to examine basal plasma oxytocin and faecal cortisol concentrations. Although outside the scope of the current paper, the hormone data is initially analysed and reported in Gnanadesikan et al. (2024).

Finally, when puppies were approximately 2 months old (mean start age = 55 days, range = 51-61 days), they participated in the DCDB over a series of three days. Puppy testing was conducted between March 2018 and April 2021. Throughout development, volunteers who raised the puppies in their homes from approximately 2-20 months of age, known as puppy raisers, also completed multiple behavioural surveys when the puppies were 6, 10, 12, and 16 months of age. The first puppies in the study turned 6 months old in July 2018 and the final puppies in the study turned 16 months old at the end of June 2022; thus, questionnaire data was collected between July 2018 and July 2022.

### Subjects

We recruited and filmed 67 litters over the enrolment period (January 2018 through February 2021). Of those, we were unable to test puppies from 5 litters born between February and mid-March 2020 on the DCDB due to COVID-19 restrictions, so these litters were dropped from the study. Of the remaining 62 litters from whom we collected video observations and puppy cognition data, we dropped 1 for accidental data loss due to hard drive malfunction and 2 for methodological inconsistencies (one litter whelped in a BC home but then spent several weeks at the CEDC prior to weaning, and the other litter was removed from the whelping pool a week earlier than all other litters in the study). In total, we completed video observations on 59 dams with litters that were included in the study (Table 1).

**Table 1.** Summary of data collection for participants included in the study.

| Category | subject | measure | n |
| --- | --- | --- | --- |
| *Rearing location* |  |  |  |
|  | dam | Private home (BC)/professional centre (CEDC) | 30/29 |
|  | puppy | Private home (BC)/professional centre (CEDC) | 118/117 |
| *Maternal behaviour observations* |  |  |  |
|  | dam | maternal behaviour score (litter level) | 59 |
|  | puppy | maternal behaviour score (individual level) | 235 |
| *Cognitive testing* |  |  |  |
|  | dam | adult DCDB scores | 58<sup>a,b</sup> |
|  | puppy | 8-week DCDB scores | 235 |
| *Raiser-completed behavioural assessments* |  |  |  |
|  | dam | 12-month C-BARQ | 33 |
|  | puppy | 6-month C-BARQ | 218 |
|  | puppy | 10-month distractibility & impulsivity | 203 |
|  | puppy | 12-month C-BARQ | 222 |
|  | puppy | 16-month distractibility & impulsivity | 201 |

<sup>a</sup> All dams were tested, but n = 58 because one dam participated as a BC dam with one litter and a CEDC dam with a subsequent litter
<sup>b</sup> For logistical reasons, 6 dams (2 BC and 4 CEDC) participated in cognition testing after the study litter was weaned

Throughout the recruitment process, we ensured that the dams in both the CEDC and BC environments were balanced on breed and parity, and that we sampled as equally as possible from among the range of options in these categories. This balancing was necessary to ensure that any differences between the two populations could be attributed to their whelping and rearing environment as opposed to other demographic factors, since past research shows that breed and parity may affect maternal style (e.g., Bray et al., 2017a). In our final sample, the average parity of CEDC dams was 2.62 and the average parity of BC dams was 2.67 (Supplementary Table S1). All dams were either full Labrador retriever, full golden retriever, or a cross between the two breeds, and so we calculated the percentage Labrador of each dam (ranging from 0—i.e., full golden—to 100—i.e., full Labrador). The average percentage Labrador for CEDC dams was 72% and the average percentage Labrador for BC dams was 66% (Supplementary Table S2). Finally, past research has also shown that litter size can affect maternal behaviour (e.g., Bray et al., 2017a). While we were unable to control this variable during the recruitment phase, once all litters were recruited, we found that this variable was also well balanced: the average size of CEDC litters was 7.4 pups per litter, while the average size of BC litters was 7.6 pups per litter (Supplementary Figure S1).

While the 59 filmed litters consisted of 439 total puppies, we subsequently selected 2-5 puppies from each litter (mean = 3.98 puppies per litter), for a total of 235 puppies, to contribute follow-up measures, including cognitive testing and raiser-completed behavioural questionnaires (Table 1). In selecting these puppies, we sought to balance sex (n = 120 F / 115 M) and prioritize puppies who would return to the Northwest and Southwest Training Centers for Canine Companions (collectively 77% of our sample) so that we could collect adult behavioural measures on as many of the dogs as possible.

### Rearing environment

Puppies were either whelped in private homes of local volunteer breeder caretakers (BC) or at the Canine Early Development Center (CEDC), a state-of-the-art facility with full-time staff dedicated to monitoring and caring for the puppies and their mothers. In both environments, puppies were kept in a whelping pool or box – to which the mother had constant access – for the first 3 weeks, and then over the 4^th^ week transitioned to living and sleeping together in a larger enclosed area. Puppies in both environments were provided with enrichment, including diverse toys, textures, and sounds. All puppies began weaning at four weeks of age, at which point a puppy kibble diet was introduced and provided three times per day. Between 7-9 weeks of age, all puppies spent at least 3 days at the CEDC and underwent veterinary exams prior to being sent to their volunteer puppy raisers, where they resided for the next ∼14-18 months. All puppies remained socially housed with their littermates up until the time they went to their individual volunteer puppy raisers.

Puppies whelped and reared with volunteer breeder caretakers lived in home environments and were cared for by the breeder caretaker(s). Depending on the home, these puppies may have been exposed to and handled by a variety of people, including children and adults, who socialized the puppies and performed basic husbandry, such as nail trims. Some puppies were also exposed to other pets, such as dogs and cats. To keep the puppies safe and healthy, Canine Companions provided husbandry and disinfection guidelines for the whelping pool or box and policies on dog interactions to minimize exposure to pathogens. Typically, puppies were closely monitored, with a human in the home at all times, for the first two weeks and then left alone with their mother for periods of time during the subsequent weeks. Breeder caretakers had access to a Canine Companions staff member via a 24/7 phone line in the event of questions or concerns about the puppies’ health, and if necessary, puppies were brought to campus to be examined by a staff veterinarian. All puppies were seen during a telemedicine appointment with Canine Companions veterinary staff at approximately 5 weeks of age. Breeder caretakers were encouraged to do early neurological stimulation (ENS) with the puppies daily from days 3-16. Puppies reared at a breeder caretaker home remained with their mother until 7-8 weeks old, when they travelled as a litter to the CEDC.

Puppies whelped and reared at the CEDC lived in a kennel environment specifically designed for dams and puppies and were cared for by CEDC staff members. The CEDC was staffed 24/7 by three shifts of employees, with a minimum of two staff members monitoring the puppies overnight and several staff members during the day. CEDC caretakers monitored the puppies both in person and via a closed-circuit video feed which transmitted and recorded footage 24/7. The CEDC has strict husbandry and disinfection policies to prevent illness, including requiring all staff to wear clean scrubs and shoes, wash their hands prior to entering the kennel area, and wear disposable gloves as appropriate. Puppies reared at the CEDC did not interact with other dogs or animals outside of their litter, although they did see and hear other litters and dams in nearby kennels. These puppies typically only interacted with CEDC caregivers and research staff, who were all adult and predominantly female. CEDC staff members socialized the puppies, performed basic husbandry such as nail trims, and did ENS exercises with the puppies daily from days 3-16. Puppies reared at the CEDC remained with their mothers until approximately 6 weeks old when they finished weaning and the dam returned to the home of her volunteer breeder caretaker. These puppies then remained with their litter until they travelled to their individual puppy raisers at 7-8 weeks old.

### Maternal behaviour observations

#### Protocol

For maternal observations in the BC environment, we helped the caretaker set up and position a provided Samsung SmartCam before the pups were two days old. This camera was zoomed in on the whelping pool and recorded continuously onto an SD card during specified periods over the first three weeks post-whelp. We later extracted 10-minute recordings of interest (without sound, described below). For maternal behaviour observations at the CEDC, we had access to dedicated camera systems from which we could automate 10-minute recordings on study litters throughout the day via Blue Iris software.

#### Scoring

All litters were video recorded in 10-minute increments for a total of 12 hours over the first three weeks—from this footage, we coded 2 hours (40 min/week) of the videos for maternal behaviour and summed the amount of time they experienced each type of behaviour. The 10-minute segments that we coded typically occurred at 4:30am, 11:00am, 4:30pm, and 11:00pm on day 3 (week 1), day 9 (week 2), and day 15 (week 3) post-birth. Deviations were rare but sometimes occurred due to technological issues or to avoid human interactions. Specifying pre-set time windows is consistent with prior literature (e.g., Foyer et al., 2016; Montgomery et al., 2025) and allowed for consistency across participants and weeks; however, we acknowledge that the trade-off is that we might have systematically underrepresented behaviours that occurred outside of those windows. Specifically, we tracked proximity (defined as a dam having all four feet in the whelping pool; the only measure that, by definition, was always identical for all littermates), licking/grooming (defined as a puppy’s fur and/or anogenital region being licked, nudged, or physically contacted by the dam’s snout), nursing (defined as a puppy suckling or attempting to suckle), and physical contact (defined as a puppy touching the dam’s body and/or face, excluding grooming and nursing). Although not further discussed in the current paper, each litter was also video recorded for an additional 24 hours each week, for a total of 72 hours over the course of the study, and this footage was specifically coded for time spent vertical nursing, a rarer behaviour defined as puppies nursing while the dam was sitting, standing, or laying on her back. Puppies wore individually identifiable collars (by colour and markings that can be differentiated in the dark via infrared light) and were marked in unique locations with small dots of paint, allowing for both daytime and nighttime observations.

### Cognitive testing

#### Protocol

All 235 puppies and 58 dams in our study participated in the Dog Cognitive Development Battery (DCDB). At the time of testing, puppies ranged from 7.3 – 8.7 weeks (mean = 7.88 weeks) and dams ranged from 1.65 – 5.18 years (mean = 3.16 years). The DCDB consists of 14 tasks presented to puppies over the course of three sessions and, with minor modifications, to adult dogs over the course of two sessions (Bray et al., 2020; Bray, Gruen, et al., 2021). The full procedures and scoring protocol for each task are detailed in Gnanadesikan et al. (2023). For the purposes of the current study, our analyses included puppy performance on all DCDB tasks and adult performance on our abbreviated DCDB battery, which includes five tasks (i.e., novel object, surprising events, human interest, the unsolvable task, and the cylinder task) relating to temperament, social tendencies, and problem solving. In the current study, 94% of puppy participants completed every single task in the DCDB, and 100% of dam participants successfully completed the five tasks of interest. However, in our subsequent analyses looking at the association between pre-pregnancy dam behaviour and maternal behaviour, we excluded six dams who were supplemented during the postnatal period with substances intended to impact mothering style (Lezama-García et al., 2019; Lyu et al., 2025; Santos, Beck, Blondel, et al., 2020)—specifically, oxytocin nasal spray and/or an ADAPTIL pheromone collar. The final sample size for each measure is reported in Table 2.

**Table 2.**
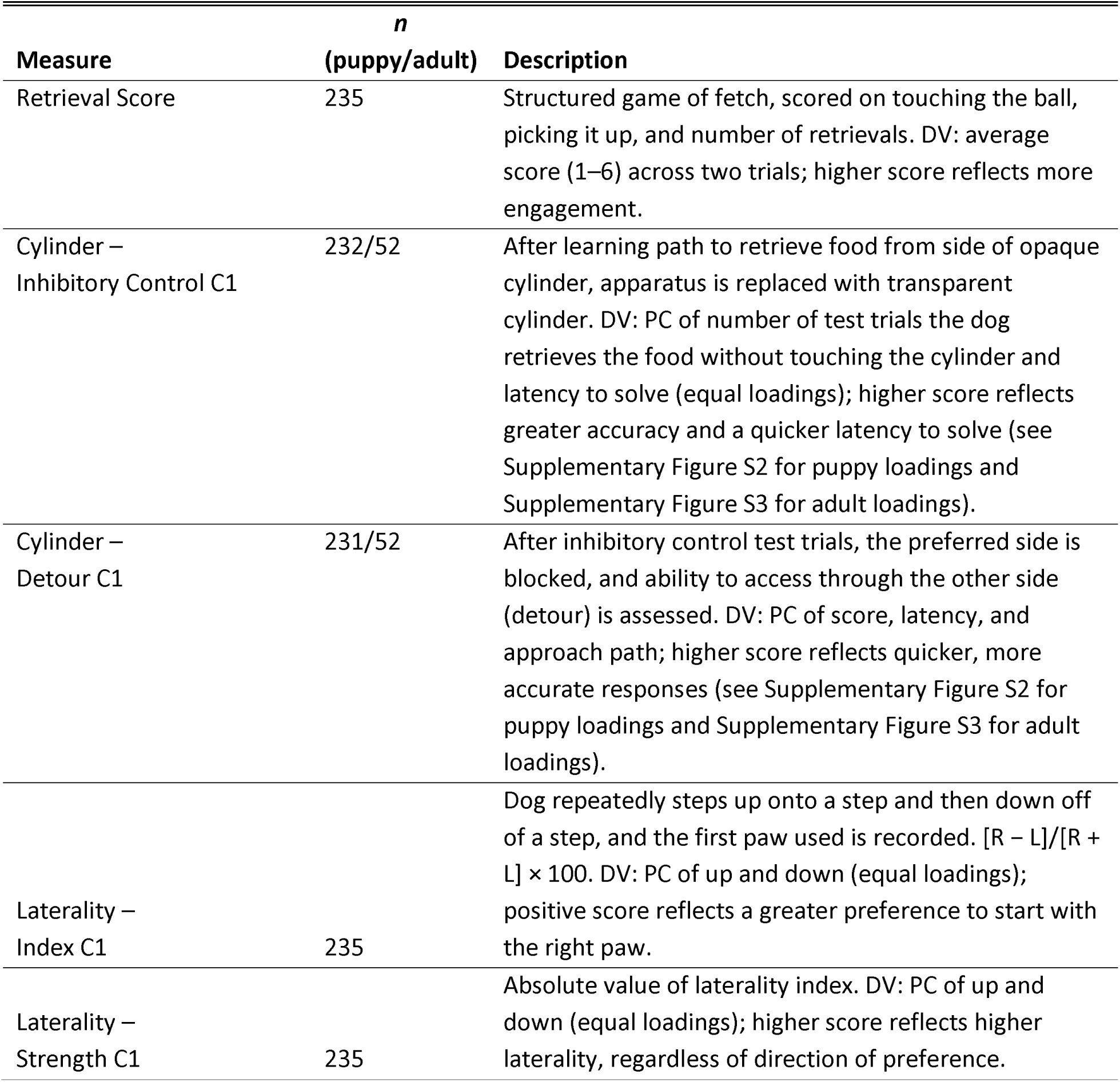
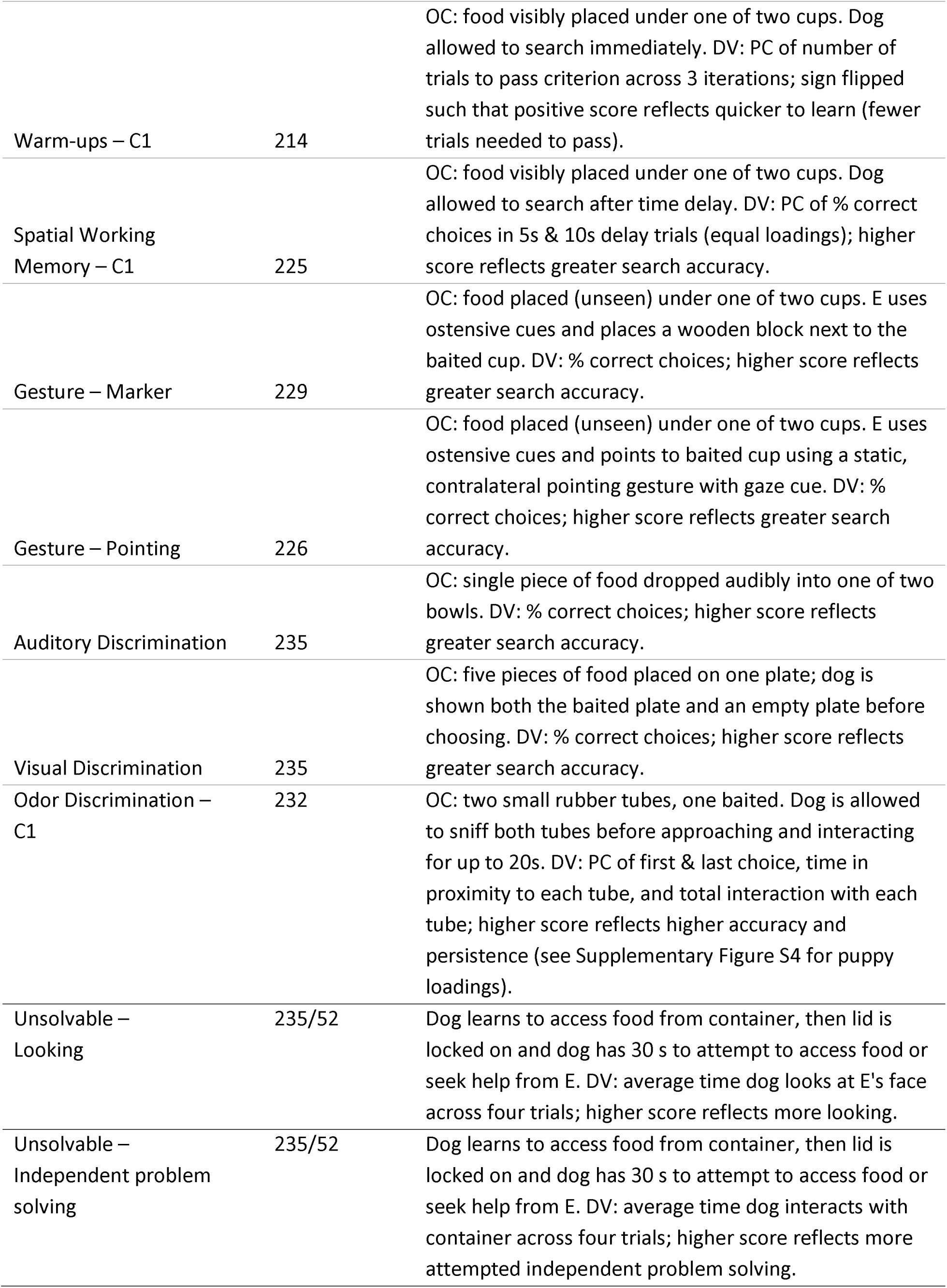
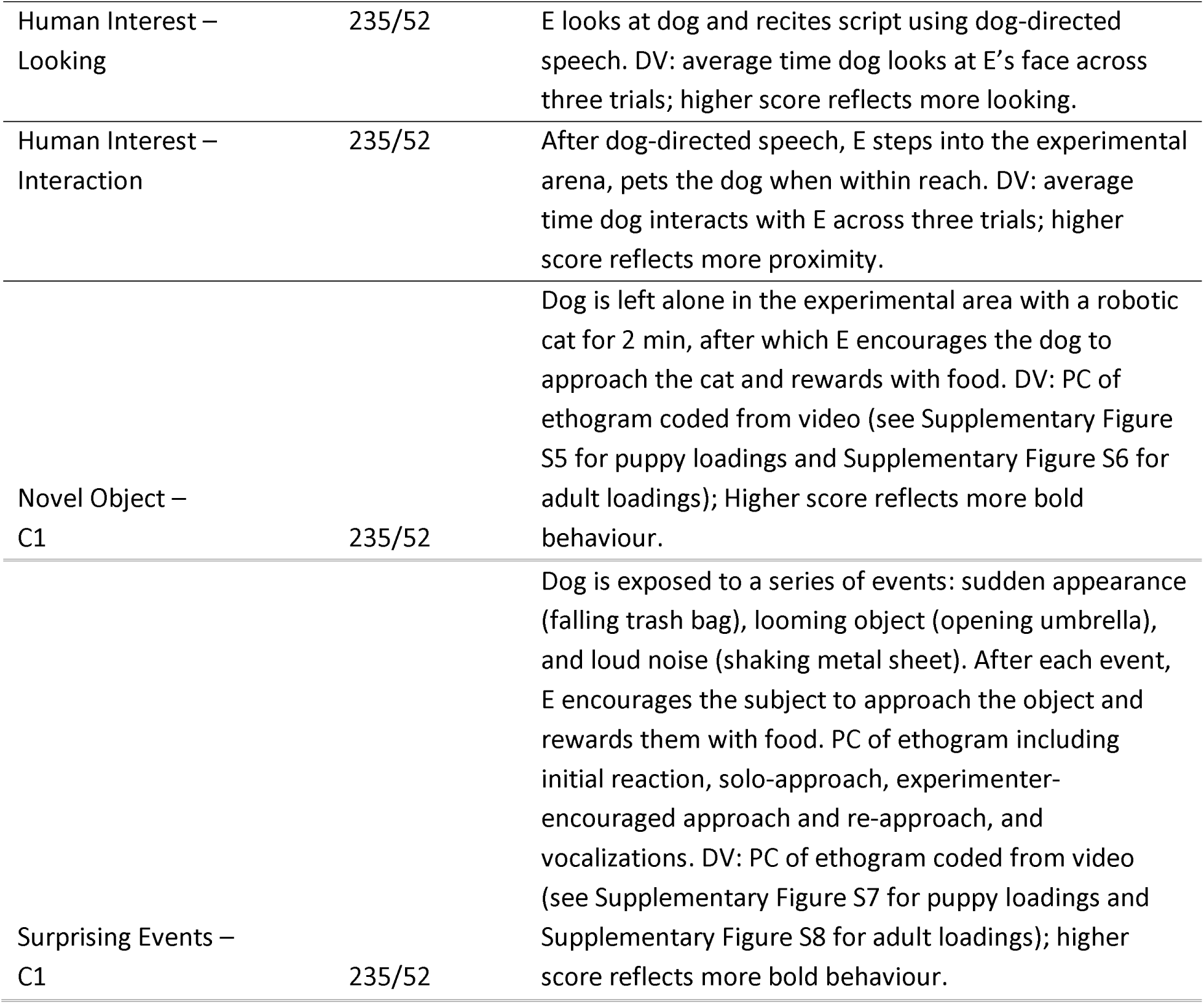
Overview of behavioural (cognitive and temperament) phenotypes and sample sizes used in subsequent analyses. OC = Object Choice; PC = Principal Component; C1 = Component 1; E = Experimenter, DV = dependent variable. Because we analysed the cylinder, unsolvable, human interest, novel object, and surprising events variables in our adult dam sample as well, those sample sizes are reported as (puppy/adult). Table adapted with permission from Gnanadesikan et al. (2024).

#### Scoring

For DCDB tasks, the number of raw outcome measures varied by task, ranging from single scores reflecting overall task performance (e.g., percent correct in object choice tasks) to many behavioural measures scored using an ethogram (e.g., novel object and surprising events tasks). For theoretical reasons and consistent with past literature (Bray, Gnanadesikan, et al., 2021), we were interested in social looking as a standalone measure, so for human interest and unsolvable we kept both outcome measures as distinct raw variables. We similarly considered the two aspects of the cylinder task – inhibitory control and reversal learning – as two separate constructs that we represented with two distinct principal components. Finally, also consistent with past literature, we considered strength and direction of laterality preference separately (Tomkins et al., 2010). For all other tasks that included more than one dependent measure, we conducted principal components analysis with all measures and retained scores on the first principal component as our outcome measure for that task. These principal components analyses were conducted on a larger dataset (puppy *n* = 468; adult *n* = 1089) associated with our ongoing work on the development of dog cognition. Component loadings are provided in the supplemental materials (Figures S2-S8) and briefly summarized in Table 2.

#### Reliability

All behavioural tasks were video recorded for reliability assessment. We obtained strong inter-rater reliability on both the puppy and dam DCDB measures, which is summarized in the supplemental materials (Supplementary Tables S3-S8).

### Raiser-completed behavioural assessments

#### C-BARQ protocol

Canine Companions sends an email (and up to 3 reminders, if necessary) to the raiser of every puppy, when the puppy turns 6 months and again when they are 12 months old, asking them to complete the Canine Behavioral Assessment & Research Questionnaire (C-BARQ, www.cbarq.org) (Serpell & Hsu, 2001). This survey consists of 101 questions related to the dog’s behaviour in ordinary settings which measure the presence and severity of broader problems like fear, aggression, and anxiety, as well as some positive traits like trainability. It takes approximately 10-15 minutes to complete. Of the 235 study puppy raisers who were sent the 6-month C-BARQ, 218 completed it, yielding a 93% response rate. When the same raisers were sent the 12-month C-BARQ, 222 completed it, yielding a 94% response rate. Additionally, 33 of the 58 dams who went on to participate in the study had a C-BARQ filled out for them when they were approximately 12 months of age, yielding a 57% response rate. Of those respondents, we excluded any dams who had been given oxytocin nasal spray and/or an ADAPTIL pheromone collar post-whelp as mentioned above, and so our sample size for analysis was 29.

#### C-BARQ scoring

For the C-BARQ measures, we used scores on the 14 primary factors (which are created by averaging scores on items in related categories), rather than raw items. Categories assessed included trainability, energy, excitability, chasing, attachment & attention seeking, separation-related problems, touch sensitivity, fear (stranger-directed, dog-directed, non-social), and aggression (stranger-directed, owner-directed, dog-directed, and familiar dog aggression). For the dams, we used scores on 11 of the 14 primary factors, excluding the following three categories from analyses due to a lack of variation: owner-directed aggression, dog-directed aggression, and familiar dog aggression. For the puppies, C-BARQ scores obtained at 6 months and 12 months were modelled separately.

#### ADHD & DIAS protocol

As part of this study, we asked raisers to provide information on participating puppies through additional surveys. At 10 months of age, we emailed puppy raisers prompting them to fill out two validated questionnaires regarding their puppy’s distractibility (Dog-ADHD questionniare; Vas et al., 2007) and impulsivity (DIAS questionnaire; Wright et al., 2011), along with demographic details. The Dog-ADHD questionnaire includes 13 questions assessing attention skills, impulsivity, and motor activity in dogs, adapted from a validated survey evaluating ADHD-related symptoms in children. The Dog Impulsivity Assessment Scale (DIAS) is an 18-item validated questionnaire assessing impulsive tendencies in dogs. The two psychometric tools were combined into one electronic survey for simplicity. Of the 235 puppies who were sent the 10-month questionnaires, 203 completed them, yielding an 86% response rate. At 16 months of age, puppy raisers were again prompted to fill out the ADHD and DIAS questionnaires, along with demographic details. 201 raisers filled out the 16-month questionnaire, again yielding an 86% response rate.

#### ADHD & DIAS scoring

In the ADHD questionnaire, each question prompts the puppy raiser to rate the frequency of a behaviour from never (0) to very often (3). The answers to six items were then averaged to form an ‘inattention’ score, while the answers to the remaining seven items were averaged to form an ‘activity-impulsivity’ score for each dog, as was done in the original paper (Vas et al., 2007). Dogs who scored highly on the ‘inattention’ score tended to be described as likely to lose interest quickly, easily distracted, having difficulty concentrating and listening, and slow to learn, especially when given complex tasks. Dogs who scored highly on the ‘activity-impulsivity’ score tended to be described as active, fidgety, and always on the move, as well as lacking self-control. In the DIAS questionnaire, the original paper found their data was best described by a three-factor solution: Behavioural Regulation, Aggression/Response to Novelty and Responsiveness (Wright et al., 2011). We found our data was best described by four components, where Aggression and Response to Novelty were distinct (Supplementary Table S9). Specifically, our four components were interpreted as Behavioural Regulation (dogs who scored high on this component were impulsive, excitable, impatient), Aggression (dogs who scored high on this component acted aggressively when excited or frustrated), Response to Novelty (dogs who scored high on this component were hesitant of and disinterested in novel things and situations), and Responsiveness (dogs who scored high on this component were easy to train, interested in new things, and agreeable). For both instruments, scores obtained at 10 and 16 months were modelled separately.

### Statistical Models

All statistical analyses were carried out in R v.4.5.1 (R Development Core Team, 2024). We used a Bayesian approach to statistical analysis. All models were fit using the brms R package (Bürkner, 2017) using an identity link function and Gaussian response distribution. Outcome and (continuous) predictor variables were processed using a rank-based inverse normal distribution and scaled and centred to have a mean of 0 and standard deviation of 1 to better meet the assumptions of linear modelling. For each model we ran 4 independent sampling chains which were merged for the posterior distribution. We used weakly regularizing priors for the beta coefficients (a normal distribution with a mean of 0 and standard deviation of 1), which is more conservative than frequentist approaches, particularly in the context of multiple hypothesis testing (Gelman & Tuerlinckx, 2000). For all models we considered the central 90% quantile range of the posterior distribution as the credible interval (CI) for a parameter, given the data and model (Kruschke, 2014).

To assess associations between maternal behaviour and offspring phenotypes, we fit multilevel models predicting the outcome variable as a function of our maternal behaviour principal component scores (hereafter, maternal behaviour scores) at a given week in development (week 1, week 2, week 3) and a set of control variables. In all models we included a random intercept for litter ID and fixed effects for dog sex, rearing location, and breed composition (percent Labrador retriever ancestry: [0,25]; (25,50]; (50,75]; (75,100]). For DCDB measures, we also included an additional covariate for age (weeks) at evaluation. Initially we attempted to also control for relatedness among subjects using an “animal model” (Wilson et al., 2010) but encountered convergence issues when also including the random intercept for litter ID. We therefore eliminated the relatedness term but retained the random effect for litter to account for non-independence of maternal effects among dogs from the same litter. To explore demographic and environmental predictors of maternal behaviour, we modelled principal components characterizing dam maternal behaviour at each week as a function of breed composition, birth season, parity, and whelping location.

## Results

### Characterizing Maternal Behaviour

As has been reported in past literature (Baqueiro-Espinosa et al., 2024; Bray et al., 2017a; Foyer et al., 2016; Guardini et al., 2017; Guardini et al., 2015; Guardini et al., 2016; Montgomery et al., 2025; Santos, Beck, Blondel, et al., 2020), we found that most mother-puppy interactions (i.e., time that the mother spends in proximity, nursing, contacting, and licking/grooming her puppies) decreased across weeks, with only vertical nursing increasing with time (Figure 2).

**Figure 2.**
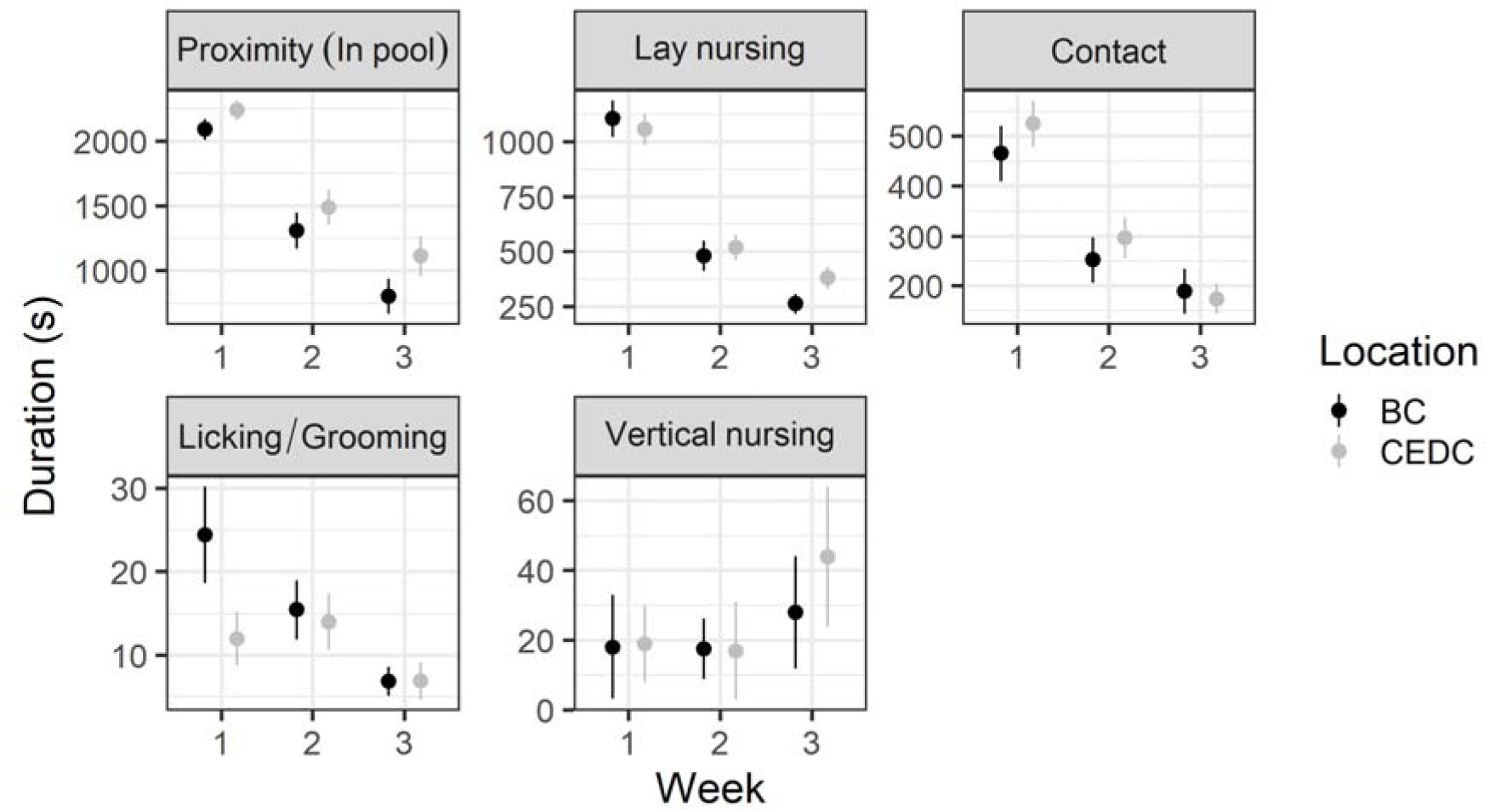
General patterns of maternal behaviour over the first three weeks post-whelp during 2400 seconds of recordings each week (raw data).

### Scoring maternal behaviour at the individual level

Along with Guardini et al. (2017; 2016), this study is among the first to individually identify the puppies and assign different scores to puppies within the same litter, based on their unique interactions with the dam. To develop summary measures for the extent of maternal interaction to which individual puppies were exposed, we performed principal component analyses. Parallel analysis of maternal behaviours directed toward individual puppies (contact, grooming, proximity, vertical nursing & lay nursing) suggested retention of a single component, and the scree plot similarly showed inflexions that were consistent with retaining a one-component solution. The presence of a single maternal behaviour component is consistent with past studies (e.g., Bray et al., 2017a; Foyer et al., 2016; Guardini et al., 2017; Guardini et al., 2016). This component explained 39% of variation in maternal care observations and was loaded positively by all measures. Variable loadings were strong for contact (0.64), grooming (0.50), proximity (0.89), and lay nursing (0.72), with a considerably weaker loading by vertical nursing (0.09), which was a rarely observed behaviour. Thus, puppies who scored high on this maternal behaviour component had mothers who were often in close proximity and who frequently interacted with their puppies via physical contact, licking, grooming, and lay nursing. These scores were used in all individual-level analyses (Figure 1).

### Scoring maternal behaviour at the litter level

For analyses of predictors of maternal behaviour, which pertain to the dam rather than the puppy as a unit of measure, we additionally generated litter-level component scores for each dam by averaging the component scores for the observed puppies from the same litter. To ensure this was an accurate representation of a dam’s behaviour, we also calculated another dam-specific maternal behaviour score by coding duration of a dam’s maternal behaviours (physical contact, licking, grooming, lay nursing, and proximity) toward all puppies in her litter (i.e., not just the study puppies) and created a second principal component score based on this data. When we compared the two dam scores (one based on averaged behaviour toward individual study pups and the other based on behaviour toward her entire litter), the correlation was extremely high (r = 0.95).

### Maternal behaviour sampling strategy

As described above, we coded and calculated maternal behaviour for all 59 litters across videos that spanned 40 minutes per week, totalling two hours per litter over the three-week observation period. However, we intentionally recorded additional video from all litters. To assess whether our primary sampling strategy of 40 minutes per week adequately captured patterns of variation derived from more extensive sampling, we conducted a methodological study using data from a subset of participants. Specifically, we coded the full amount of data collected (240 minutes per week, totalling 12 hours per litter over the three-week observation period) for 10 dams and their 40 study puppies over three weeks, half of which were whelped and reared in the CEDC environment and half of which were whelped and reared in the BC environment. We generated principal component scores using this full dataset and compared these scores to those used in the main analyses (i.e., scores created using only 40 minutes per week of data, totalling 2 hours per litter, from those same dogs). We found that scores derived from the full dataset correlated strongly with scores derived from the primary dataset at both the puppy (r = 0.80; Figure S9) and litter (r = 0.84; Figure S10) level. We thus conclude that two hours of observations over three weeks is sufficient to capture variation in maternal care.

### Can we predict the maternal behaviour that a dam will display towards her pups?

To determine if dam behavioural characteristics, measured pre-pregnancy, predict subsequent maternal care, we fit separate models for each behaviour, while controlling for breed composition, birth season, parity, and rearing location (Figure 3, Supplementary Table S10). Our only finding from our DCDB predictors was that dams who exhibited the most cognitive flexibility in the cylinder detour task (i.e., achieved high accuracy and were quick to solve) showed higher levels of maternal behaviour at week 1 post-whelp (β_Week 1_= 0.18; 90% CI = 0.061, 0.29) and, although the credible interval did include zero, a similar pattern was observed in week 3 (β_Week 3_= 0.18; 90% CI = −0.006, 0.36). No associations were found between dams’ responses on the other tasks (unsolvable, human interest, surprising events, and novel object) and their later maternal behaviour.

**Figure 3.**
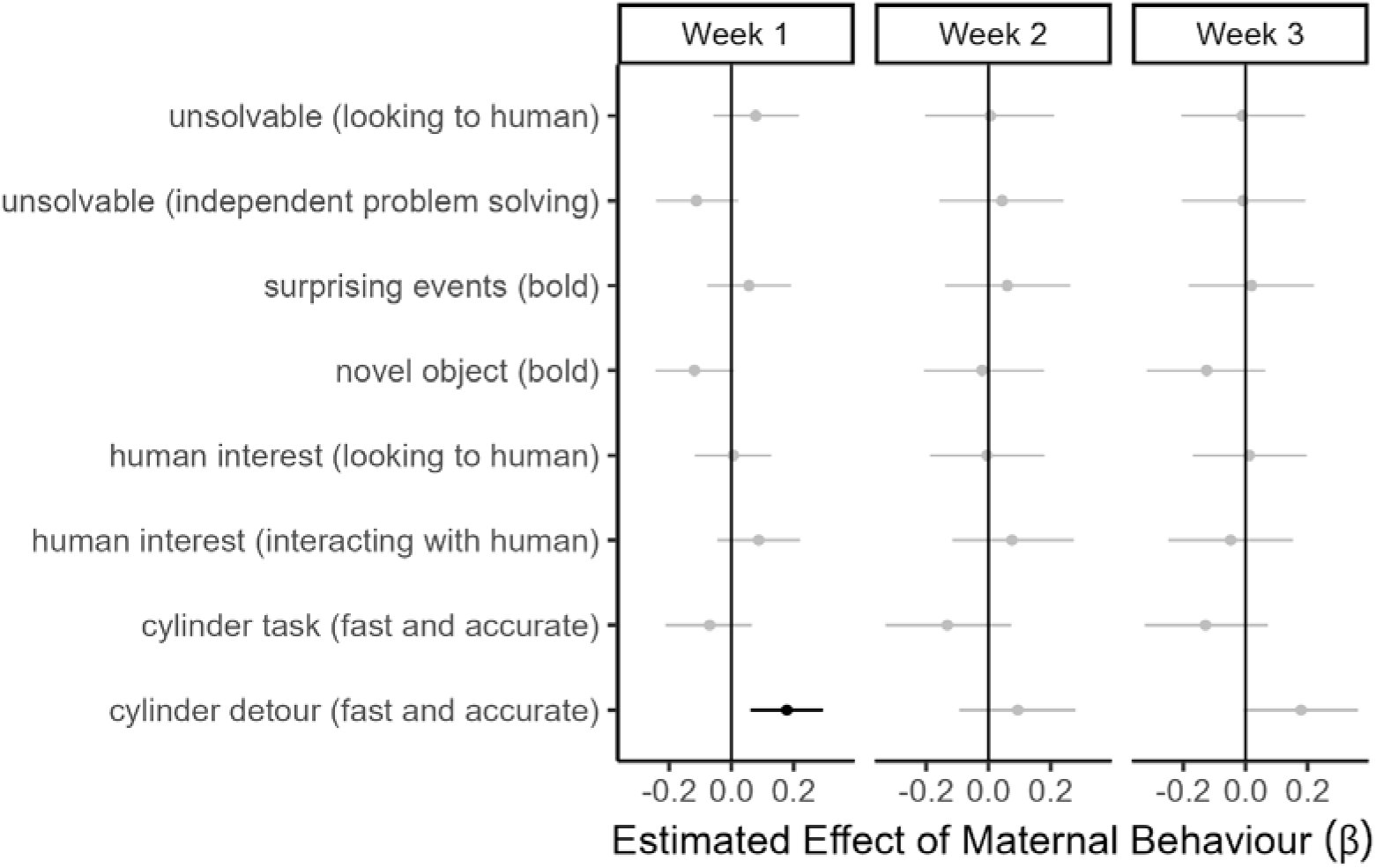
Dam DCDB measures predicting maternal behaviour.

In contrast, we found several associations between the 12-month C-BARQ scores for dams, provided by puppy raisers prior to the dog being chosen as a breeder, and their subsequent maternal behaviour as observed in the study (Figure 4, Supplementary Table S11). Specifically, dams who were rated lower on the excitability factor (“displaying strong reactions to potentially exciting or arousing events…and has difficulty settling down after such events”) displayed higher levels of maternal behaviour in the first (β_Week1_ = −0.17; 90% CI = −0.33, −0.01) and second weeks (β_Week2_= −0.28; 90% CI = −0.55, −0.002) post-whelp. Dams who were rated lower on chasing (“chases cats, birds, and/or other small animals, given the opportunity”) also displayed higher levels of maternal behaviour in the second week (β_Week2_ = −0.46; 90% CI = −0.74, −0.17) post-whelp. Finally, dams who were rated higher on trainability (“shows willingness to attend to the owner, obeys simple commands, learns quickly, fetches objects, responds positively to correction, and ignores distracting stimuli”) displayed higher levels of maternal behaviour in the second week (β_Week2_= 0.33; 90% CI = 0.04, 0.61) post-whelp.

**Figure 4.**
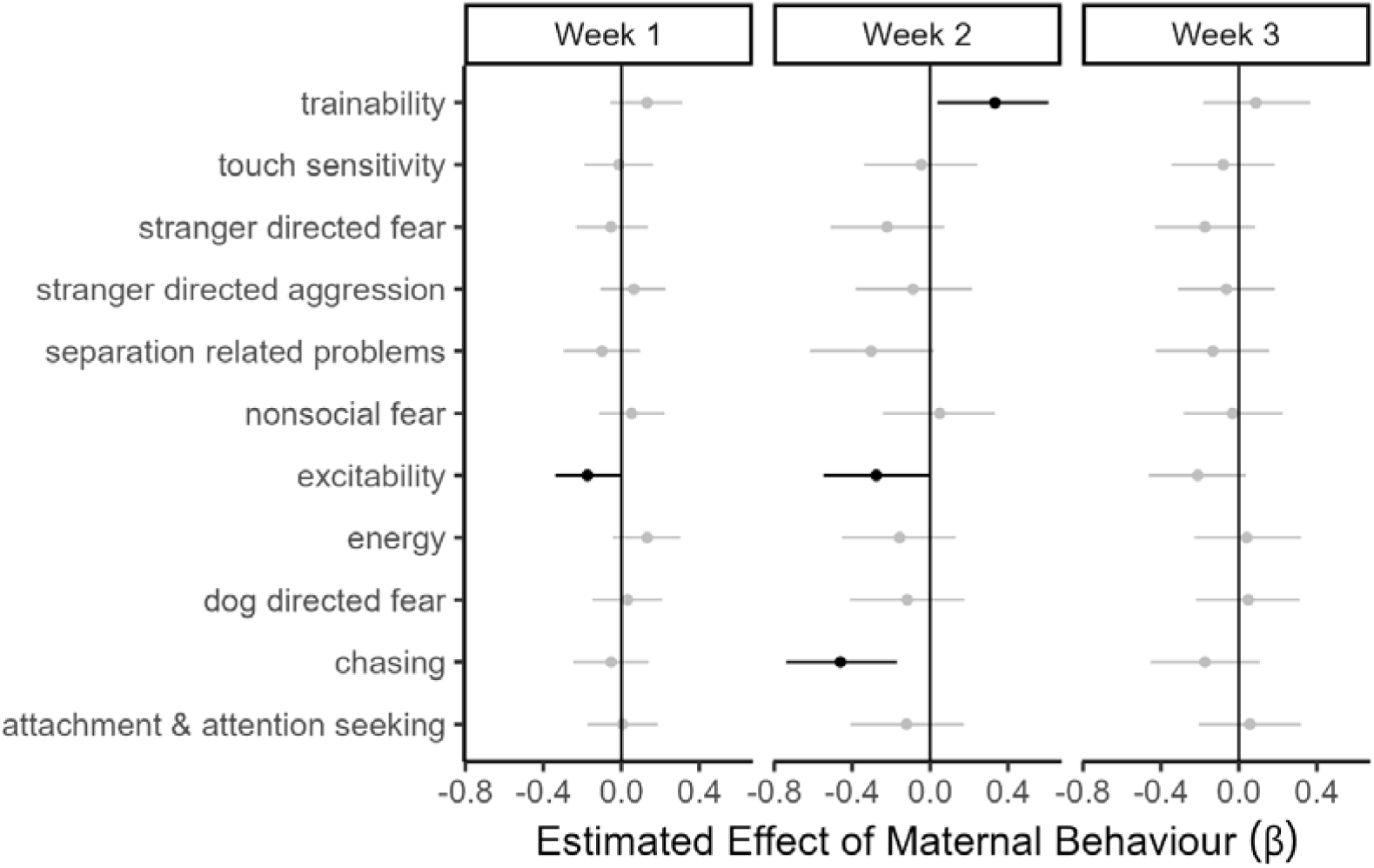
Dam 12-month C-BARQ scores predicting subsequent maternal behaviour.

### Does early rearing environment affect maternal behaviour?

To assess demographic and environmental factors that might affect maternal behaviour, we fit a linear mixed model predicting maternal behaviour scores as a function of fixed effects for week of life (1-3), litter size (3-11), dam parity (1-5), breed composition (0-25% Labrador, 26-50% Labrador, 51-75% Labrador, 76-100% Labrador), birth season (January-March, April-June, July-September, October-December), and rearing location (BC or CEDC). As we saw with the raw score patterns, maternal behaviour decreased across the three-week period following parturition, with the largest decrease between weeks 1 and 2 (β_week 2_ = −1.03; 90% CI = −1.22, −0.85; β_week 3_= −1.55; 90% CI = −1.73, −1.36). Neither parity, litter size, breed composition, nor rearing location was associated with maternal behaviour scores (Supplementary Table S12). However, maternal behaviour scores did vary by season, with the highest scores observed during the fall and winter months and the lowest during the late spring and summer (estimates relative to April-June: β_January-March_ = 0.39; 90% CI = 0.05, 0.73; β_July-September_ = 0.19; 90% CI = −0.14, 0.54; β_October-December_ = 0.47; 90% CI = 0.14, 0.81). The intraclass-correlation coefficient for this model was 0.35, suggesting moderate week-to-week consistency of individual differences in maternal behaviour.

### Does early rearing location affect puppy behaviour?

To determine if early rearing location affected puppy behaviour, we first measured behavioural outcomes of puppies via puppy behavioural testing around 8 weeks of age. The descriptive statistics of measures collected during these testing sessions for all puppies are summarized in Table 3. As reported in our previous research (Bray et al., 2020), puppies performed above chance on all tasks where they were presented with two choices (Supplementary Table S13); this suggests that—on average—puppies at this age are cognitively capable of using a variety of cues to make informed choices. The only exception was the odour control task, where puppies performed at chance, just like adult dogs (Bray, Gruen, et al., 2021), indicating that they were unable to solve the problem based on odour cues and strengthening the evidence that puppies are successfully using the provided cues in the other object choice tasks.

**Table 3.** Puppy performance on the DCDB: Descriptive statistics.

| *variable* | *n* | *mean* | *sd* | *minimum* | *maximum* | *standard error* |
| --- | --- | --- | --- | --- | --- | --- |
| age at testing (days) | 235 | 55.13 | 2.50 | 51.00 | 61 | 0.16 |
| parity | 235 | 2.64 | 1.45 | 1.00 | 5 | 0.10 |
| birth order | 235 | 4.63 | 2.44 | 1.00 | 11 | 0.16 |
| warmups 1: trials to criterion | 214 | 7.85 | 4.87 | 4.00 | 25 | 0.33 |
| warmups 2: trials to criterion | 231 | 6.57 | 4.25 | 4.00 | 23 | 0.28 |
| warmups 3: trials to criterion | 229 | 6.14 | 3.64 | 4.00 | 26 | 0.24 |
| retrieval: average score | 235 | 3.60 | 1.11 | 1.00 | 5 | 0.07 |
| retrieval: tally | 235 | 3.74 | 3.86 | 0.00 | 19 | 0.25 |
| laterality: bias strength | 235 | 44.39 | 29.20 | 0.00 | 100 | 1.90 |
| laterality: laterality index | 235 | -5.19 | 52.96 | -100.00 | 100 | 3.45 |
| human interest: avg look time | 235 | 5.93 | 4.11 | 0.00 | 19.64 | 0.27 |
| human interest: avg interaction time | 235 | 18.73 | 8.13 | 0.00 | 27.86 | 0.53 |
| cylinder: familiarization score | 232 | 8.06 | 3.52 | 4.00 | 23 | 0.23 |
| cylinder: inhibitory control score | 232 | 39.60 | 22.44 | 0.00 | 100 | 1.47 |
| cylinder: reversal score | 231 | 19.81 | 20.85 | 0.00 | 87.5 | 1.37 |
| unsolvable task: avg time looking at human (s) | 235 | 1.07 | 1.22 | 0.00 | 7.34 | 0.08 |
| arm pointing | 226 | 64.12 | 18.39 | 8.33 | 100 | 1.22 |
| communicative marker | 229 | 69.87 | 19.12 | 8.33 | 100 | 1.26 |
| odour control trials | 224 | 49.16 | 16.82 | 0.00 | 87.5 | 1.12 |
| memory (5s) | 227 | 75.33 | 19.73 | 16.67 | 100 | 1.31 |
| memory (10s) | 225 | 67.93 | 20.76 | 0.00 | 100 | 1.38 |
| memory (15s) | 146 | 65.75 | 20.51 | 16.67 | 100 | 1.70 |
| memory (20s) | 85 | 63.53 | 22.05 | 16.67 | 100 | 2.39 |
| visual discrimination | 234 | 91.88 | 12.07 | 50.00 | 100 | 0.79 |
| auditory discrimination | 234 | 61.06 | 17.79 | 12.50 | 100 | 1.16 |
| odour discrimination:<br>first choice | 232 | 56.82 | 19.19 | 0.00 | 100 | 1.26 |
| odour discrimination:<br>final choice | 232 | 70.76 | 19.95 | 16.67 | 100 | 1.31 |

Whereas maternal behaviour did not differ by rearing location, we did find differences in puppy performance on certain DCDB tasks based on whether the puppies were whelped and reared in a volunteer breeder caretaker home or the CEDC (Figure 5). Specifically, CEDC puppies generally interacted more during the play breaks of the human interest task (β = 0.63; 90% CI = 0.36, 0.90), were more accurate and quicker to solve during the cylinder inhibitory control trials (β = 0.34; 90% CI = 0.08, 0.59), achieved higher accuracy on the visual discrimination task (β = 0.29; 90% CI = 0.05, 0.53), and were more strongly lateralized when stepping up and down (β = 0.27; 90% CI = 0.03, 0.51). CEDC puppies were also less accurate on the pointing task (β = −0.40; 90% CI = −0.68, −0.11) and demonstrated less cognitive flexibility and were slower to solve during the cylinder detour trials (β = −0.25; 90% CI = −0.48, −0.02). During the surprising events task, CEDC puppies were less bold, more reactive and slower to recover, and more vocal (β = −0.37; 90% CI = −0.64, −0.10). Finally, although the credible interval did include zero, a similar pattern towards being less bold and more vocal was observed in the CEDC puppies on the novel object task as well (β = −0.21; 90% CI = −0.43, 0.007).Thus, across DCDB measures, some aspects of puppy performance was associated with rearing location, but in different directions across tasks. These findings raise the question of which traits are ultimately most important for training outcomes, along with the possibility of whether there might be interventions that can be implemented in either BC homes or the CEDC environment to address shortcomings.

**Figure 5.**
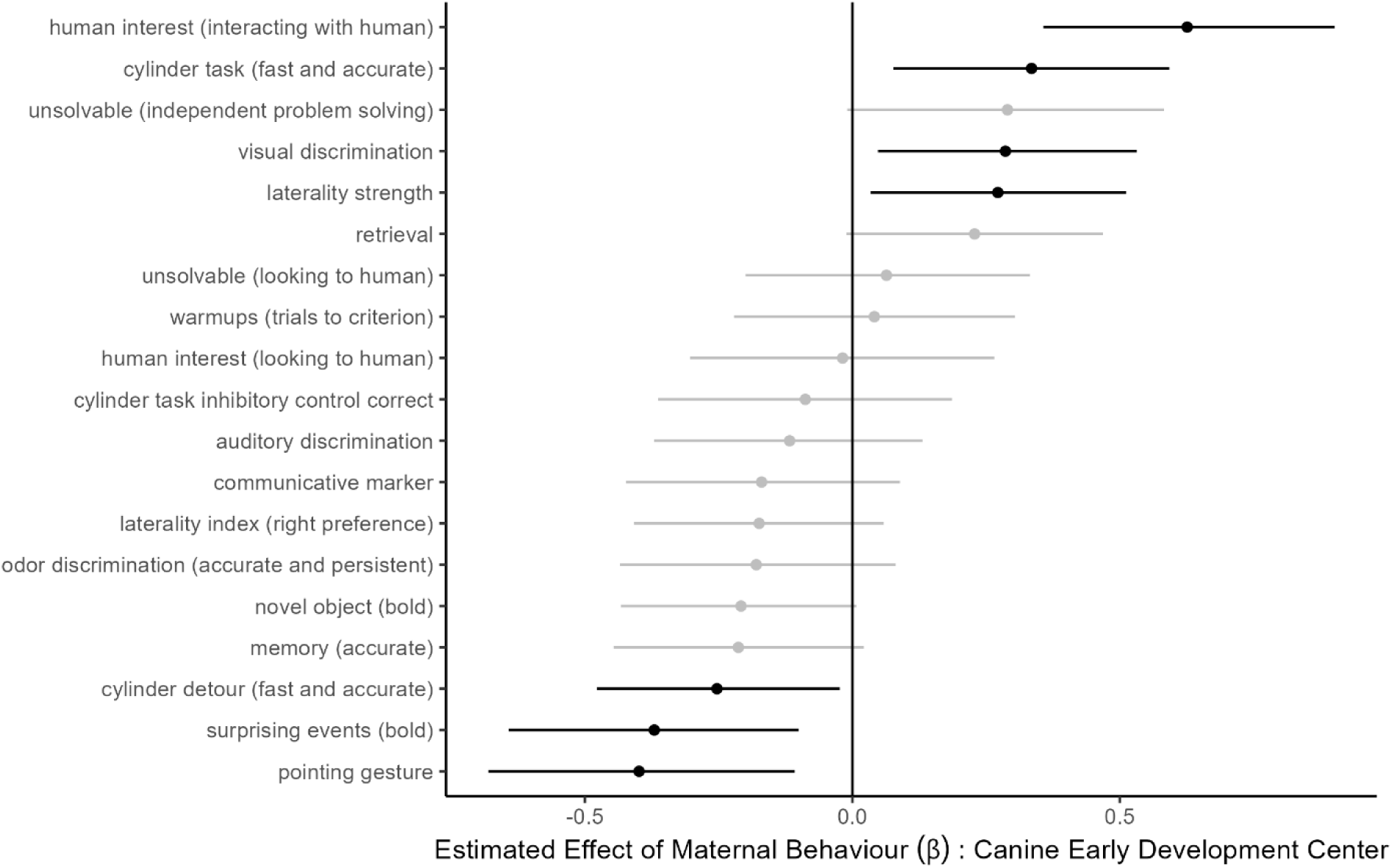
Association between rearing location and DCDB measures. The reference level is BC-reared puppies, and the plot shows the effects for CEDC-reared puppies.

### How is maternal behaviour associated with puppy behaviour?

#### Behaviour assessed via DCDB

We observed seven associations between maternal behaviour and offspring performance on the DCDB at ∼8 weeks of age for which the credible interval did not include 0 (Figure 6; Supplementary Table S14). Most of these associations related to maternal behaviour appeared during week 2 (five associated measures), with fewer associations during week 1 (two associated measures), and no associations during week 3. Specifically, exposure to higher levels of maternal behaviour during the first week of life was associated with better performance on the cylinder task (β = 0.17; 90% CI = 0.04, 0.29), a measure of executive function, but with worse performance on the auditory discrimination task (β = −0.14; 90% CI = −0.27, −0.01). Exposure to higher levels of maternal behaviour in the second week of life was associated with poorer outcomes on several DCDB measures, including a higher number of trials required to meet criterion in a warm-up procedure (β = −0.16; 90% CI = −0.29, −0.03), less accurate responses during the communicative marker task (β = −0.13; 90% CI = −0.26, −0.002), and worse performance in both the olfactory (β = −0.13; 90% CI = −0.25, −0.01) and auditory discrimination (β = −0.16; 90% CI = −0.28, −0.04) tasks. We also observed a negative association between maternal care in week 2 and a right limb bias on the motor laterality task (β = −0.16; 90% CI = −0.28, −0.04); in other words, as maternal care increased, puppies were more left-biased. In sum, the maternal style experienced by puppies during the second week of life had the most associations with DCDB performance at 8 weeks of age.

**Figure 6.**
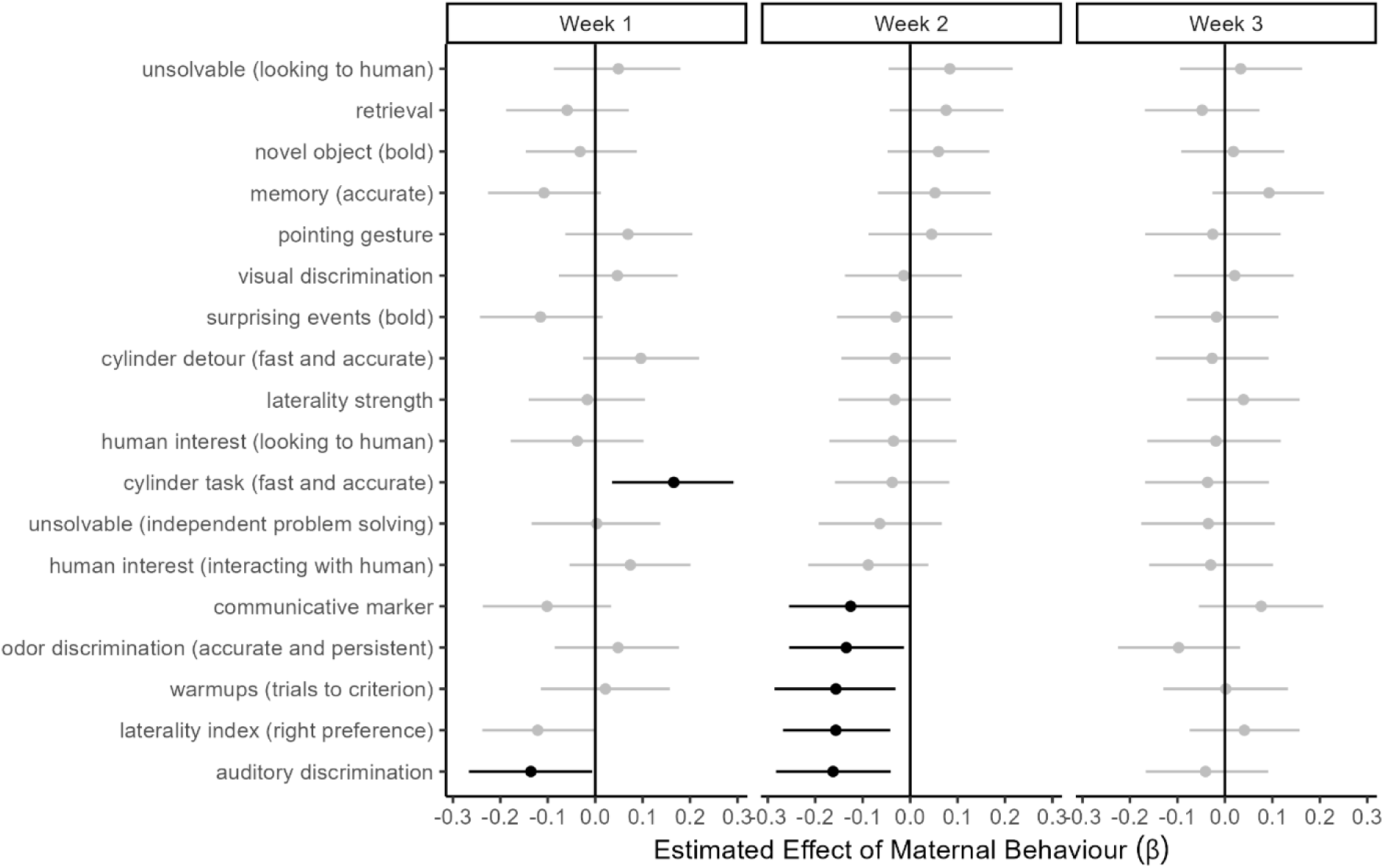
Associations between maternal behaviour and 8-week-old puppy DCDB performance. Associations for which the credible interval does not include 0 are shown in black.

#### Behaviour assessed via C-BARQ

Maternal care was associated with several behavioural measures from the C-BARQ at 6 and 12 months of age (Figure 7, Supplementary Table S15). Dogs who experienced higher levels of maternal behaviour during the first week of life exhibited more problematic behaviours related to non-social fear (at 6 months: β = 0.14; 90% CI = 0.004, 0.27; “fearful or wary responses to sudden or loud noises, traffic, and unfamiliar objects and situations”) and chasing (at 12 months: β = 0.14; 90% CI = 0.002, 0.28). Increased maternal behaviour in week 2 was associated with more problematic separation-related behaviour (at both 6 months: β = 0.14; 90% CI = 0.02, 0.26; and 12 months: β = 0.13; 90% CI = 0.003, 0.25; “vocalizes and/or is destructive when separated from the owner”) and dog-directed fear (6 months: β = 0.14; 90% CI = 0.009, 0.27), but also with fewer problems involving stranger-directed fear (6 months: β = −0.15; 90% CI = −0.27, −0.03). Lastly, dogs who experienced more maternal behaviour during week 3 exhibited more problems with owner-directed aggression (at 6 months: β = 0.14; 90% CI = 0.03, 0.26), lower energy levels (at 12 months: β = −0.12; 90% CI = −0.25, −0.009; “energetic, ‘always on the go’, and/or playful”), and, although the credible interval did include zero, a similar pattern of more problematic separation-related behaviour (at 12 months: β = 0.12; 90% CI = −0.005, 0.24). Although we found associations for each week, maternal care experienced by puppies in the second week of life again seems especially important, as it exhibited the most associations, and especially an enduring association with separation-related behaviour at both 6 and 12 months.

**Figure 7.**
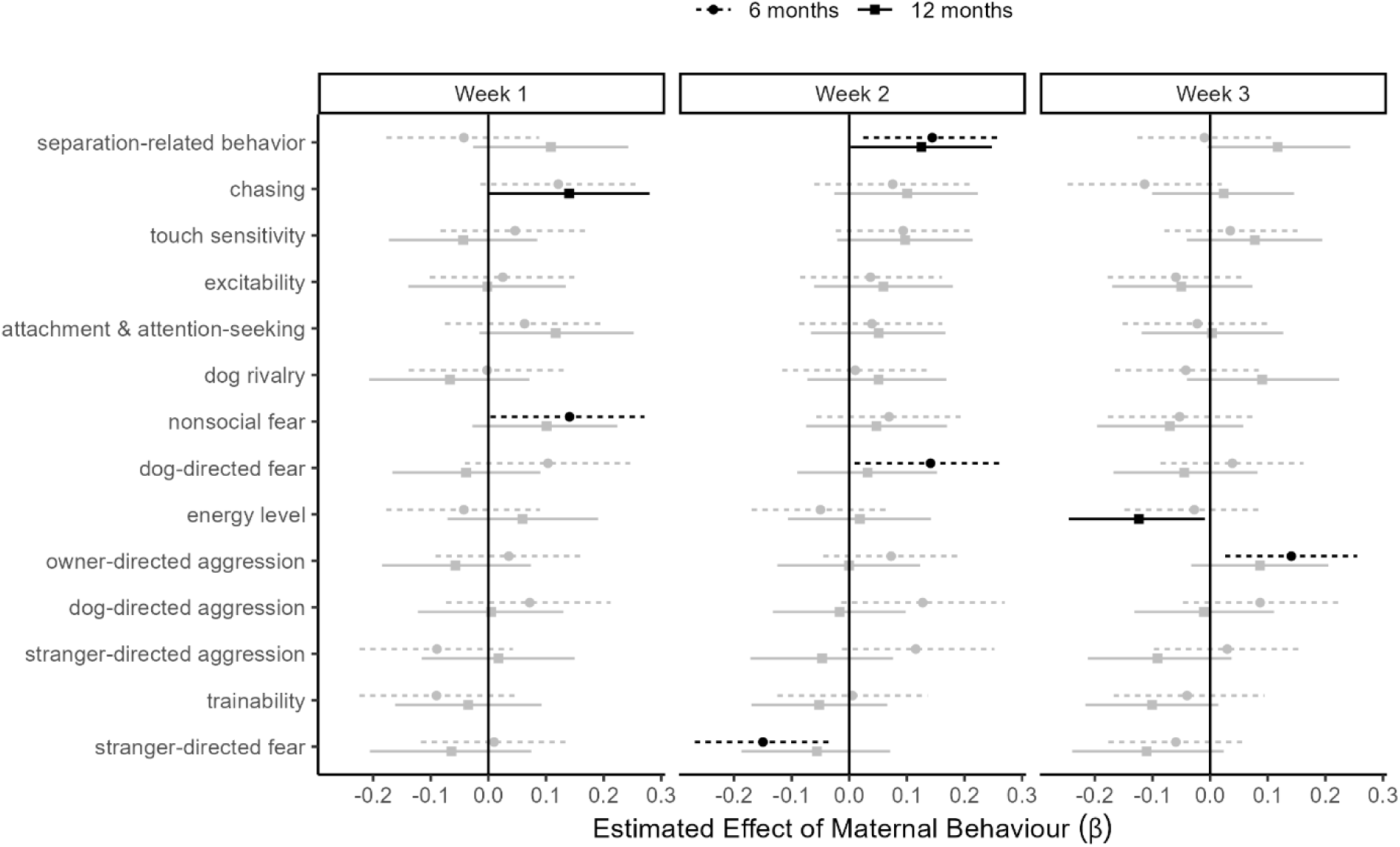
Associations between maternal behaviour and offspring C-BARQ scores at 6 months and 12 months of age. Associations for which the credible interval does not span across zero are shown in black.

#### Behaviour assessed via the Dog Impulsivity Assessment Scale (DIAS) & the Dog-ADHD Rating Scale

Finally, we found several associations between maternal care and behavioural measures from the DIAS and Dog-ADHD surveys at 10 and 16 months of age (Figure 8, Supplementary Table S16). In fact, the activity & impulsivity measure was the only one with no association to maternal behaviour. Specifically, dogs exposed to more maternal care during the first week of life scored lower on responses to novelty (DIAS) at 10 months (β = −0.14; 90% CI = −0.27, −0.01) and 16 months (β = −0.17; 90% CI = −0.30, −0.04) of age, meaning they were more comfortable around and interested in novel things and situations. Maternal care in week 2 was positively associated with higher aggression (DIAS) at 10 months of age (β = 0.14; 90% CI = 0.02, 0.26) and poorer behavioural regulation (DIAS) at 16 months of age (β = 0.13; 90% CI = 0.00, 0.25). Lastly, dogs exposed to higher levels of maternal care during week 3 scored higher on inattention (Dog-ADHD) at 10 months of age (β = 0.14; 90% CI = 0.01, 0.26), and lower on responsiveness (DIAS) at 16 months of age (β = −0.19; 90% CI = −0.32, −0.07).

**Figure 8.**
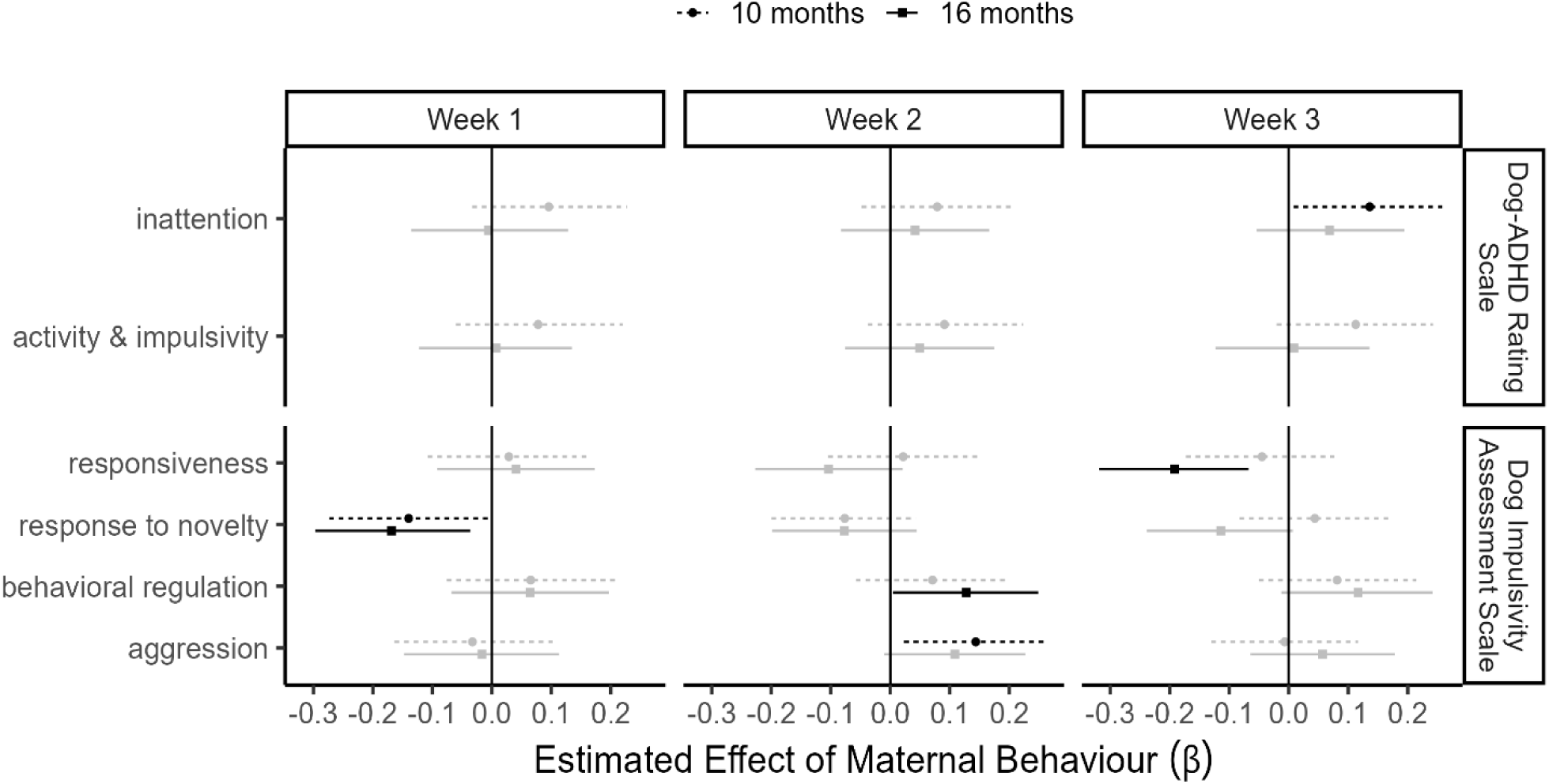
Associations between maternal behaviour and offspring DIAS and Dog-ADHD Rating Scale response scores at 10 and 16 months of age. Associations for which the credible interval does not span across zero are shown in black.

## Discussion

In this study, we collected cognitive and behavioural data from a sample of 59 dams and 235 puppies to explore the predictors and downstream effects of variation in maternal care and the early rearing environment. Our characterization of maternal care was largely consistent with previous literature in that we video recorded and tracked typical maternal behaviours – proximity, contact, nursing, and licking/grooming – to describe the maternal care displayed by each dam (Santos, Beck, & Fontbonne, 2020). Using our data, we addressed three orienting questions: First, can dam behaviour assessed pre-pregnancy predict subsequent maternal care? Second, how does early rearing environment affect maternal care and subsequent puppy characteristics? And third, what are the effects of maternal care on puppy cognitive and behavioural phenotypes, from 8 weeks through 16 months? Below, we address key findings with respect to each of these questions.

The potential to identify a dam’s proclivity to display certain levels of maternal care is beneficial to working dog organizations and pet dog breeders alike. Past research in guide (Bray et al., 2017b) and detection (Montgomery et al., 2025) dogs has demonstrated a link between maternal behaviour and subsequent puppy outcome in those respective programs; thus, knowing what to look for in a potential breeding dog could pay dividends in bolstering success rates within working dog programs. Likewise, as we continue to learn more about how maternal care fosters various behavioural phenotypes, private pet dog breeders could be more selective by choosing breeding dams who will produce fewer puppies displaying the types of behavioural problems that often lead to relinquishment (Bray, Otto, et al., 2021; Czerwinski et al., 2016).

To assess potential behavioural predictors of maternal care in the current study, we looked at dams’ pre-pregnancy responses to our abbreviated DCDB tasks. In general, we found those tasks were not useful predictors of subsequent maternal behaviour. The only association we found was between quick and accurate performance on the cylinder detour task (indicating cognitive flexibility) and higher levels of maternal care in the first week post-whelp (Figure 3).

We also looked at the C-BARQ scores of dams whose puppy raisers had completed the questionnaire when they were approximately 12 months old, which only applied to approximately half of our sample (*n* = 29). We found that dogs who were reported to display less excitability, more trainability, and lower levels of chasing behaviour tended to provide more maternal care (Figure 4). It seems reasonable that more excitable dams may be more likely to move around rather than lie with their puppies, as reflected in lower levels of care and a tendency to engage in more vertical nursing. Similarly, since dams are often confined to a separate area with their neonates, those dogs who engaged in more human-oriented separation behaviours pre-pregnancy may be more prone to similarly display those behaviours while separated from their human caregivers with their litters, possibly resulting in distraction from mothering behaviours. Montgomery et al. (2025) also recently looked at associations between C-BARQ scores and maternal behaviour and found dams who scored high on excitability spent more time nursing in the vertical position. In the sense that vertical nursing has been found to be inversely related to other aspects of maternal care (e.g., as reported in the current study, overall levels of maternal care decline across the first three weeks while vertical nursing increases), this result is consistent with our own. Furthermore, Montgomery et al. (2025) report that higher scores for human-oriented separation-related behaviour predict lower maternal care. We observed this same pattern across all three weeks in our data as well, although all the credible intervals included zero (Figure 4). Together with Montgomery et al. (2025), our results suggest that the C-BARQ could be a helpful tool in evaluating future dams, and it appears to be more predictive of future maternal behaviour than the DCDB. From a practical standpoint, the C-BARQ is quite feasible for working dog organizations to implement, allowing for the assessment of hundreds of dogs while requiring minimal investment of staff time and resources (Hare et al., 2024).

We found a strong effect of time post-whelp on maternal behaviour: specifically, the amount of maternal care steadily decreased over time, as has been consistently reported in the literature (Santos, Beck, & Fontbonne, 2020). We also found that dams who whelped their litter in warmer months (April through June) displayed less maternal behaviour than dams who whelped in colder months (October to March). One possible reason for this difference is that the ambient temperature in the room during October through March might be colder, leading the dam and puppies to spend more time close to each other in the whelping box. The daylight hours are also shorter in those months, affecting both temperature and light in ways that might lead to dogs and caretakers spending more time inside and with the puppies. Other research (Baqueiro-Espinosa et al., 2022; Foyer et al., 2016) has not found birth season to influence maternal care; however, Foyer et al. (2016; 2013) did find an effect of birth season on puppy behaviour during a temperament test and Baqueiro-Espinosa et al. (2022) found that dams experienced more difficult whelps during the winter.

No other demographic factors were associated with variation in maternal care, and specifically, we did not find an effect of litter size. While two previous studies (Baqueiro-Espinosa et al., 2024; Bray et al., 2017a) reported that aspects of maternal behaviour decreased with increasing litter size, two other recent studies (Montgomery et al., 2025; Santos, Beck, Blondel, et al., 2020) did not find an effect of litter size on maternal care. It is possible that the different findings between the studies are related to breed composition. Like Montgomery et al. (2025), the population of working dogs in our study consisted of only retrievers. Surprisingly, we also found that parity did not make a difference. While this finding mirrors that of Foyer et al. (2016), it differs from our past study in guide dogs that found that the more experienced the mother, the less maternal behaviour she displayed (Bray et al., 2017a). While we also found no effect of breed composition, this finding can likely be attributed to the fact that the breeds used were all retrievers and thus highly similar.

We did not observe a difference in maternal care displayed by dams whelping and rearing their puppies in a professional breeding facility (the CEDC) as compared to those in a home environment (Figure 2). To our knowledge, our study is the first to directly compare maternal behaviour across different whelping and rearing locations. The two other studies that tangentially address this topic did find differences in behaviour by rearing environment but also used different methodologies. Tiira & Lohi (2015) reported that dogs who whelped at their permanent homes took better care of their offspring compared to dogs who travelled to another location to whelp and rear their litters. However, rather than directly observing the dogs, this study queried owners via questionnaires, relying on their memories and perceptions of the dam’s maternal care when visiting the breeder and/or following up with the breeder. Baquiero-Espinosa and colleagues (2022) compared the behaviour of dams who had been reared in the breeding kennel to dams who had been born elsewhere and arrived at the breeding kennel later in life. Dams who were born and reared in the breeding kennel had significantly easier whelps and spent more time nursing their puppies during the first 24 hours postpartum compared to dams who were born elsewhere. Our finding that maternal behaviour in this population does not appear to be related to whelping and rearing location may be of practical importance for working dog organizations who utilize one or both types of locations for their breeding programs.

Next, we looked at associations between behaviour and the early rearing location that puppies inhabited during early life. We found an association between rearing location and several aspects of puppy behaviour measured with the DCDB at 8 weeks of age, consistent with prior research documenting the influence of rearing environment on puppy behaviour development (Figure 5; Appleby et al., 2002; Lenkei et al., 2019; Majecka et al., 2020). For example, compared to BC puppies, CEDC puppies showed better performance on the cylinder task test trials, meant to measure motor inhibition, but inferior performance on the cylinder task detour trials, meant to measure cognitive flexibility. We also found that, compared to BC puppies, CEDC puppies were more lateralized on a behavioural task measuring paw preference. We initially included the behavioural lateralization task in our battery because laterality is theorized to reflect cerebral lateralization and has previously been associated with cognitive (e.g., problem-solving skills; Magat & Brown, 2009) and temperament (e.g., confidence in the face of novelty; Batt et al., 2009) traits. However, in the current population we did not find that it correlated with any of these constructs as operationalized by the DCDB (Gnanadesikan et al., in press). Given that finding, we are not sure why there is a difference in laterality strength between CEDC and BC puppies, nor the implications. This finding is nonetheless interesting, given that laterality strength on a similar measure in guide dogs was found to be significantly associated with the ultimate success of a dog in graduating from the training program (Tomkins et al., 2012).

Our strongest findings were that puppies born and reared at the CEDC spent more time in proximity to the experimenter during the play breaks of the human interest task and were less skilled at following a communicative pointing gesture. It is possible these results are related to the differing types of human interaction and exposure that these puppies experienced, leading to differences in social and communicative behaviour. Though all puppies in the study had daily interaction with people, puppies reared in a BC home may have spent more time interacting with the same human(s), whereas puppies reared at the CEDC likely interacted with a wider variety of humans and spent less time interacting with any given person. Lenkei et al. (2019) also found differences in human social interaction in 8-week-old puppies raised in a home environment compared to those raised in a kennel. Puppies raised in a home environment maintained gaze with the experimenter longer than those raised in a kennel. Compared to puppies raised in a home, puppies raised in a kennel spent more time close to an experimenter sitting silently in the pen, though they spent less time interacting with the experimenter during a recall test. While there is not a clear pattern of behaviour related to human social interaction, as Lenkei and colleagues (2019) note, these findings suggest that developmental differences related to social and communicative interactions with humans may emerge at an early age in puppies and reinforce the significance of early socialization with humans.

We also found that puppies raised at the CEDC were less confident and more likely to vocalize during a surprising events temperament task (Figure 3). This difference might be attributed to the environmental surroundings in the CEDC compared to a home environment. Though diligent care is taken to expose the puppies in the CEDC to a variety of stimuli, a home environment may have a greater variety of sights, sounds, and smells and would have been more unpredictable than the CEDC environment. These early differences in experiences may have influenced the puppies’ confidence levels when exposed to novel objects and surprising events during the temperament tasks. Majecka et al. (2020) found that puppies reared indoors in a home were more likely to receive scores characterizing them as confident, lacking aggressive tendencies, and able to cope with novelty than puppies raised in an outdoor kennel. Appleby et al. (2002) found that a home-based early rearing environment was associated with a reduced probability of avoidance behaviour and aggression toward unfamiliar people, and reduced aggression during a veterinary exam later in life. Our findings contribute to the literature on the effect of early rearing environment on puppy behaviour development and suggest that working dog organizations and breeders who house puppies in a kennel environment may want to take thoughtful care to socialize the puppies to humans and expose them to a broad variety of stimuli. It is important to note that all DCDB puppy testing was completed at the CEDC the week of the scheduled veterinary checks, prior to going to puppy raiser homes. As a result, BC puppies travelled to be tested and at the time of testing were being temporarily housed in an environment which was novel to them. Thus, we cannot confidently dissociate effects related to early rearing location from experiences more proximate to DCDB testing.

Next, separate from rearing location, we found that the maternal care experienced by the puppies was related to their later behaviour as measured by the DCDB, an experimental behavioural assay, and as reported through caretaker questionnaires. We found seven associations between maternal care and performance on the DCDB, with Week 2 emerging as the timepoint with the most associations. Most of those associations indicated that more maternal care was generally related to less desirable outcomes (See Figure 6). Intriguingly, associations between pre-pregnancy dam behaviour and subsequent maternal behaviour were also most evident in Week 2, therefore making it the week that was most predictable for maternal behaviour as well as most predictive of offspring behaviour on the DCDB. The maternal behaviour rodent literature similarly finds there is a critical time window—specifically, the first week post-birth—where maternal interactions are particularly important for shaping later-life offspring outcomes (Caldji et al., 1998; Liu et al., 1997; Weaver et al., 2004). Based on the current study and others (Bray et al., 2017b), the critical period in dogs appears to begin about one week later, which is likely reflective of the reality that the postnatal development of dogs progresses more slowly than that of rodents.

Past literature has found associations between levels of maternal care and performance on arena, isolation, and novel object tests in 8-week-old puppies (Baqueiro-Espinosa et al., 2025; Guardini et al., 2016, 2017). Though we did not find associations in the current study between maternal behaviour and the DCDB temperament tasks, including novel object and surprising events, considered together with previous research our results indicate that the level of maternal care in the first three weeks has a measurable impact on 8-week-old puppy behaviour as evaluated by behaviour tests. It remains to be seen if performance at 8 weeks on certain DCDB tasks is indicative of future adult behaviour and success in the program, and if so, which traits are most important. For example, on the DCDB, we found that dogs who experienced less maternal care showed a right limb bias. This finding is consistent with past literature, which separately reports that both a right paw preference (Tomkins et al., 2012) and lower levels of maternal care (Bray et al., 2017b) are linked to greater guide dog success.

Our study also provides evidence that the association between maternal care on puppy behaviour endures beyond the first 8 weeks of life. Specifically, we found several associations between maternal care and behaviours reported on caretaker questionnaires at 6, 10, 12 and 16 months of age.

Dogs who experienced more maternal care displayed less inhibition around humans, as measured by an increase in owner-directed aggression and a decrease in stranger-directed fear reported on the C-BARQ at 6 months of age. Similarly, the DIAS results also revealed a positive link between maternal behaviour and offspring aggression before 12 months of age, although this association was not observed at later ages. These findings are consistent with research by Foyer et al. (2016) who found that German shepherd military working dogs who experienced higher levels of maternal care were more likely to engage in social behaviour with humans and had higher scores on aggression at age 15-18 months.

Higher levels of maternal care were also associated with human-oriented separation-related behaviours reported on the C-BARQ at 6 and 12 months of age. Considered together with our finding that maternal care is negatively associated with some aspects of inhibition around humans and the findings of Foyer et al. (2016), it seems possible that the dam-puppy relationship dynamic may exert an influence on the later puppy-human attachment dynamic. Guardini and colleagues (2017) also found a relationship between maternal care and interest in an unfamiliar person and human-oriented separation-related behaviours in 8-week old puppies. They suggest that this could be related to the attachment bond between the dam and her offspring and that seeking security from humans is a valuable trait for dogs who are separated from their litters early in life. It is possible that for puppies who experience a higher level of maternal care, separation from the mother may be more difficult. These puppies may benefit by being less inhibited and more comfortable around humans, allowing them to more readily seek support and build relationships with human partners.

Interestingly, the behavioural disinhibition appears specific to social situations with humans, as puppies who experienced more maternal care scored higher on C-BARQ measures of non-social fear and dog-directed fear at 6 months of age (though these associations disappeared by the 12-month evaluation point). It has been suggested that the quantity of maternal care may help to mediate stress responses and help puppies to adapt to their environments (Guardini et al., 2015), which has been demonstrated in rats (Czerwinski et al., 2016). Tiira and Lohi (2015) found that fearful behaviour and anxieties were linked to poor maternal care during puppyhood using a questionnaire administered to dog owners. In contrast, other research (Bray et al., 2017b; Guardini et al., 2017) has found a positive association between maternal care and stress-related behaviours. Evidence points to a systemic effect of maternal care on fear and anxiety-related behaviours in dogs; however, the direction of the effect requires further research and may be based on the type of instrument used to measure the behaviour, the age at evaluation, and the range of variation present.

In addition to the relationships with aggression, fear, and separation-related behaviours, we found associations between maternal care and other DIAS measures potentially considered undesirable for service dogs. Dams providing more maternal care produced puppies who scored higher on inattention at 10 months of age, higher on behavioural regulation (i.e., were more impulsive, excitable and impatient) at 16 months of age, and lower on responsiveness (i.e., were harder to train) at 16 months of age.

Our prior work within guide dogs also found high levels of maternal behaviour were associated with some potentially unsuitable behaviours and undesirable outcomes in young adulthood. Specifically, puppies who received more maternal care displayed higher activity levels while in isolation, were quicker to vocalize at a novel object, had more difficulty with problem-solving tasks, and were less likely to graduate as guide dogs (Bray et al., 2017b). While there are similarities in how maternal behaviour appears to be impacting later offspring behaviour between the two datasets, it remains to be seen how maternal care will impact eventual program outcomes within the current population of service dogs. Successful service and guide dogs need to display some common behavioural traits, such as trainability, confidence, and a lack of body sensitivity (Amirhosseini et al., 2025; Bray, Otto, et al., 2021; Duffy & Serpell, 2012). Yet they are distinct roles and the literature suggests there are also unique qualities required for success in each role (Bray, Otto, et al., 2021). Thus, a future direction of our research is to examine how maternal behaviour relates to offspring adult behaviour, measured via the DCDB, and eventual outcome in the service dog program specifically.

Since we have limited knowledge about the causes and consequences of variation in maternal care, we estimated associations between maternal behaviour and a wide range of variables related to the dam, her rearing environment, and characteristics of her progeny. Given the highly exploratory nature of this work, testing the robustness of these associations, through replication in independent samples, presents an important priority of future research.

Our results provide insight into potential early environmental influences on several aspects of dog behaviour, which are relevant to both working and companion animals. Some of the effects seem to go in opposite directions in terms of desirability; thus, whether these traits are deemed desirable or undesirable may depend on the working role and the way the traits are managed. For example, dogs that experienced higher levels of maternal care exhibited an increased interest in human interaction and were also more susceptible to separation-related behavioural problems as young adults, perhaps stemming from attachment dynamics during the first weeks of life (Dietz et al., 2019). Additionally, a higher level of maternal care was associated with less inhibition around humans. Though these traits may be undesirable at the extremes, they may ultimately be qualities of a successful service dog in that they represent the strong desire to form a relationship with a human. In contrast, behaviours not appropriate for service work may be appropriate for other roles. For example, Foyer et al. (2016)found that increased maternal care was associated with aggression in young adult dogs, which is a behaviour considered desirable for military working dogs performing detection and protection roles (Bray, Otto, et al., 2021). Furthermore, several of the behavioural associations with higher levels of maternal care that we report, including measures of aggression, fear, and inattention, were not observed beyond 12 months of age on the current assessments. Thus, our future work will explore whether these behaviours persist into adulthood and ultimately affect success as a service dog.

Whereas environmental influences on common dog behaviour problems are widely recognized, few studies have considered the consequences of experiences during the first weeks of life. Our results suggest that this period may contribute importantly to the aetiology of diverse behavioural outcomes and be an important consideration when choosing breeding dams for working roles (Czerwinski et al., 2016; Foyer et al., 2016).

Collectively, our work builds on existing knowledge regarding the formative roles of early experiences. A deeper understanding of the predictors and effects of maternal behaviour will allow for more effective selection of breeders in dog populations, both working and companion, leading to optimal early rearing conditions that enhance dog welfare and support successful human-animal relationships.

## Supporting information

Raw Data

## Data Availability

Data are available as electronic supplementary material.

## Notes

### Competing Interest Statement

The authors have declared no competing interest.

### Summary of Updates

Version 2 reflects the peer-reviewed and accepted manuscript in Animal Behaviour. The manuscript and supplemental files now include additional robustness analyses requested during peer review.

### Funding

- American Kennel Club Canine Health Foundation: 02518
- Office of Naval Research: N00014-17-1-2380, N00014-20-1-2545
- U.S. National Science Foundation: DGE-1746060

