## Supplementary Material for "Maternal care and early rearing environment influence puppy behaviour and cognition"

This Supplemental Information accompanies the peer-reviewed and accepted version of the manuscript.

*Supplementary Material*

*Cognitive Testing Reliability.* All behavioural tasks were video recorded for reliability assessment. Additionally, while most behavioural variables were coded in real time, the following tasks were later coded from video: novel object and surprising events, as well as select variables from cylinder (latency during inhibitory control and reversal learning trials and first side correct during reversal learning trials), unsolvable (average time manipulating object) and odor discrimination (time at right and left elbow, from which the variables persistence, time at correct response, and time at incorrect response were subsequently calculated).

For data coded in real time, independent coders watched the video of all trials for 20% of randomly selected subjects, and we calculated interrater reliability using Pearson correlation for continuous variables and Cohen’s κ for categorical variables. For all measures that were coded from video, two coders independently scored the data for any given task. The primary coder scored all data for analysis, and the second coder scored all trials for 20% of randomly selected subjects to allow us to assess reliability. These analyses were carried out in R v.4.5.1 (R Development Core Team, 2024).

Reliability was excellent for all live-coded data; see Supplementary Table S3 (Cohen's *κ*: mean = 0.93; Pearson's *r*: mean = 0.94) for the inter-rater agreement of all puppy live-coded cognitive measures. Reliability was also strong for the video-coded data; Supplementary Table S4 reports inter-rater agreement for the video-coded measures of puppy cognitive task performance (Cohen's *κ* = 0.98; Pearson's *r*: mean = 0.97), Supplementary Table S5  reports inter-rater agreement for raw items on the novel object task (Cohen's *κ*: mean = 0.86; Pearson's *r*: mean = 0.98), and Supplementary Table S6  reports inter-rater agreement for raw items on the surprising events task, including reactions to a sudden appearance, looming object, and loud noise (Cohen's *κ*: mean = 0.95; Pearson's *r*: mean = 0.95).

Reliability was also excellent for the live-coded dam measures. This data included the two measures of human interest, average looking time (Pearson's *r* = 0.96) and average interaction time (Pearson's *r* = 0.97). Similarly, reliability was strong for the video-coded dam measures. Supplementary Table S7 reports inter-rater agreement for dam performance on the novel object task (Cohen's *κ*: mean = 0.93; Pearson's *r*: mean = 0.96), and Supplementary Table S8 reports inter-rater agreement for dam performance on the surprising events task (Cohen's *κ*: mean = 0.84; Pearson's *r*: mean = 0.89).

**Table S1**

Breakdown of study participants by parity and whelping location.

| Number of parturitions | CEDC dam | BC dam | TOTAL dams |
| --- | --- | --- | --- |
| 1 | 9 | 9 | 18 |
| 2 | 6 | 6 | 12 |
| 3 | 5 | 7 | 12 |
| 4 | 5 | 2 | 7 |
| 5 | 4 | 6 | 10 |

**Table S2**

Breakdown of study participants by breed and whelping location.

| Breed (% Labrador) | CEDC dam | BC dam | TOTAL dams |
| --- | --- | --- | --- |
| 0-25% | 5 | 7 | 12 |
| 26-50% | 5 | 3 | 8 |
| 51-75% | 0 | 3 | 3 |
| 76-100% | 19 | 17 | 36 |
| Average % Labrador | 71.79 | 65.77 |  |

**Table S3**

Reliability statistics (Cohen’s κ, Pearson’s r) for puppy live-coded measures in the Dog Cognitive Development Battery (DCDB)

| Variable | *N* | Statistic | Value |
| --- | --- | --- | --- |
| Retrieval: average score | 36 | *r* | 1.00 |
| Retrieval: tally count | 36 | *r* | 1.00 |
| Laterality | 36 | κ | 0.98 |
| Human interest: avg. look time | 36 | *r* | 0.93 |
| Human interest: avg. interact time | 36 | *r* | 1.00 |
| Cylinder: inhibitory control score | 35 | κ | 0.88 |
| Cylinder: reversal learning score | 35 | κ | 0.74 |
| Unsolvable task: avg. time looking at human | 41 | *r* | 0.84 |
| Communicative marker | 36 | κ | 0.96 |
| Arm pointing | 36 | κ | 1.00 |
| Odor control trials | 36 | κ | 0.98 |
| Memory | 41 | κ | 0.97 |
| Visual discrimination | 36 | κ | 0.97 |
| Auditory discrimination | 36 | κ | 0.96 |
| Odor discrimination: first choice | 36 | κ | 0.88 |
| Odor discrimination: final choice | 36 | κ | 0.91 |

**Table S4**

Reliability statistics (Cohen’s κ, Pearson’s r) for puppy video-coded measures of the cognitive tasks in the Dog Cognitive Development Battery (DCDB).

| Variable | *N* | Statistic | Value |
| --- | --- | --- | --- |
| Cylinder: latency (inhibitory control trials) | 35 | *r* | 0.98 |
| Cylinder: latency (reversal learning trials) | 35 | *r* | 1.00 |
| Cylinder: first side correct (reversal learning trials) | 35 | κ | 0.98 |
| Unsolvable task: avg. time manipulating object | 41 | *r* | 0.98 |
| Odor discrimination: time at right elbow | 36 | *r* | 0.95 |
| Odor discrimination: time at left elbow | 36 | *r* | 0.96 |

**Table S5**

Reliability statistics (Cohen’s κ, Pearson’s r) for puppy measures in the novel object task.

| Variable | *N* | Statistic | Value |
| --- | --- | --- | --- |
| Alone: time to first approach | 41 | *r* | 1.00 |
| Alone: proximity | 41 | *r* | 1.00 |
| Alone: no. of approaches | 41 | κ | 0.90 |
| Alone: contact | 41 | *r* | 1.00 |
| Alone: orient | 41 | *r* | 0.98 |
| Alone: time to first vocalize | 41 | *r* | 0.88 |
| Social 1: time to approach | 41 | *r* | 1.00 |
| Social 1: approach score | 41 | κ | 0.94 |
| Social 2: time to approach | 41 | *r* | 1.00 |
| Social 2: approach path | 41 | κ | 0.75 |

**Table S6**

Reliability statistics (Cohen’s κ, Pearson’s r) for puppy measures in the surprising events task.

| Variable | *N* | Statistic | Value |
| --- | --- | --- | --- |
| Sudden appearance: initial reaction | 41 | *r* | 0.85 |
| Sudden appearance: time to first approach | 41 | *r* | 0.99 |
| Sudden appearance: initial approach path | 41 | κ | 0.96 |
| Sudden appearance: time to first contact | 41 | *r* | 1.00 |
| Sudden appearance: initial contact | 41 | *r* | 0.98 |
| Sudden appearance: time to re-approach | 41 | *r* | 0.94 |
| Sudden appearance: re-approach path | 41 | *r* | 0.89 |
| Looming object: initial reaction | 41 | *r* | 0.89 |
| Looming object: time to first approach | 41 | *r* | 0.99 |
| Looming object: initial approach path | 41 | κ | 0.96 |
| Looming object: time to first contact | 41 | *r* | 0.98 |
| Looming object: initial contact | 41 | *r* | 0.91 |
| Looming object: re-approach path | 41 | κ | 0.97 |
| Loud noise: initial reaction | 41 | *r* | 0.80 |
| Loud noise: time to first approach | 41 | *r* | 1.00 |
| Loud noise: initial approach path | 41 | κ | 0.97 |
| Loud noise: time to first contact | 41 | *r* | 0.99 |
| Loud noise: initial contact | 41 | *r* | 0.98 |
| Loud noise: vocal intensity | 41 | κ | 0.91 |
| Loud noise: time to re-approach | 41 | *r* | 1.00 |
| Loud noise: re-approach path | 41 | κ | 0.99 |
| Alone: orient to experimenters | 41 | *r* | 0.94 |
| Alone: time to first vocalize | 41 | *r* | 0.99 |
| Alone: vocal intensity | 41 | κ | 0.90 |

**Table S7**

Reliability statistics (Cohen’s κ, Pearson’s r) for dam measures from the novel object task.

| Variable | *N* | Statistic | Value |
| --- | --- | --- | --- |
| Alone: time to first approach | 35 | *r* | 1.00 |
| Alone: proximity | 35 | *r* | 0.98 |
| Alone: no. of approaches | 35 | κ | 0.89 |
| Alone: contact | 35 | *r* | 1.00 |
| Alone: orient | 35 | *r* | 0.99 |
| Alone: time to first vocalize | 35 | *r* | 0.80 |
| Social 1: time to approach | 35 | *r* | 0.99 |
| Social 1: approach score | 35 | κ | 0.94 |
| Social 2: time to approach | 35 | *r* | 0.99 |
| Social 2: approach path | 35 | κ | 0.96 |

**Table S8**

Reliability statistics (Cohen’s κ, Pearson’s r) for dam measures in the surprising events task.

| Variable | *N* | Statistic | Value |
| --- | --- | --- | --- |
| Sudden appearance: initial reaction | 41 | *r* | 0.81 |
| Sudden appearance: time to first approach | 41 | *r* | 1.00 |
| Sudden appearance: initial approach path | 41 | κ | 0.91 |
| Sudden appearance: initial contact | 41 | *r* | 0.99 |
| Sudden appearance: time to re-approach | 41 | *r* | 0.84 |
| Sudden appearance: re-approach path | 41 | κ | 0.62 |
| Looming object: initial reaction | 41 | *r* | 0.75 |
| Looming object: time to first approach | 41 | *r* | 0.91 |
| Looming object: initial approach path | 41 | κ | 0.87 |
| Looming object: time to first contact | 41 | *r* | 0.92 |
| Looming object: initial contact | 41 | *r* | 0.96 |
| Looming object: time to re-approach | 41 | *r* | 0.83 |
| Looming object: re-approach path | 41 | κ | 0.93 |
| Loud noise: initial reaction | 41 | *r* | 0.78 |
| Loud noise: time to first approach | 41 | *r* | 1.00 |
| Loud noise: initial approach path | 41 | κ | 0.79 |
| Loud noise: time to first contact | 41 | *r* | 1.00 |
| Loud noise: initial contact | 41 | *r* | 0.98 |
| Loud noise: time to re-approach | 41 | *r* | 0.74 |
| Loud noise: re-approach path | 41 | κ | 0.94 |
| Alone: orient to experimenters | 41 | *r* | 0.78 |

**Table S9**

Components and loadings of the PCA over the items from the Dog Impulsivity Assessment Scale (DIAS) questionnaire. Items that loading strongly (>|0.40|) on any given component are bolded. The eigenanalysis of the correlation matrix of the items indicated that the first, second, third, and fourth principal components accounted for 22%, 10%, 9%, and 8% of the total variance, respectively.

| **DIAS item** | **Component 1: Behavioral Regulation** | **Component 2:**  **Aggression** | **Component 3: Response to Novelty** | **Component 4: Responsiveness** |
| --- | --- | --- | --- | --- |
| Extreme physical signs when excited | **0.47** | 0.23 | 0.23 | 0.36 |
| Excitement can lead to fixed repetitive behavior | **0.46** | 0.21 | 0.19 | 0.40 |
| Dog is considered to be very impulsive | **0.79** | 0.11 | -0.06 | 0.06 |
| Dog doesn’t like to be approached or hugged | 0.23 | 0.10 | 0.33 | 0.25 |
| Dog becomes aggressive when excited | 0.11 | **0.85** | 0.00 | 0.00 |
| Dog appears to be ‘sorry’ after it has done something wrong | -0.12 | 0.10 | 0.07 | **0.51** |
| Dog does not think before it acts | **0.64** | 0.11 | -0.11 | -0.12 |
| Dog can be very persistent | **0.68** | 0.17 | -0.04 | -0.03 |
| Dog may become aggressive if frustrated with something | 0.14 | **0.84** | 0.00 | 0.00 |
| Dog is easy to train (reverse scored) | **0.42** | 0.10 | 0.20 | **-0.41** |
| Dog is not keen to go into new situations (reverse scored) | -0.16 | 0.01 | **-0.71** | -0.21 |
| Dog takes a long time to lose interest in new things (reversed scored) | 0.02 | 0.10 | 0.13 | **-0.51** |
| Dog calms down very quickly after being excited (reverse scored) | **0.69** | 0.00 | 0.11 | -0.07 |
| Dog appears to have a lot of control over how it responds (reverse scored) | **0.69** | -0.01 | 0.20 | -0.23 |
| Dog is very interested in new things and new places (reverse scored) | -0.01 | -0.04 | **0.76** | -0.23 |
| Dog reacts very quickly | 0.18 | -0.03 | **-0.53** | 0.39 |
| Dog is not very patient | **0.61** | -0.04 | -0.06 | 0.07 |
| Dog seems to get excited for no reason | **0.56** | 0.31 | 0.06 | 0.22 |

**Table S10.** Associations between dam cognitive and behavioral scores on the abbreviated DCDB and subsequent maternal behaviour. CI = Credible Interval.

|  | Week 1 | |  | Week 2 | |  | Week 3 | |
| --- | --- | --- | --- | --- | --- | --- | --- | --- |
| **Dependent Measure** | **Beta** | **90% CI** |  | **Beta** | **90% CI** |  | **Beta** | **90% CI** |
| novel object (bold) | -0.12 | -0.24, 0.008 |  | -0.02 | -0.21, 0.18 |  | -0.12 | -0.32, 0.06 |
| surprising events (bold) | 0.06 | -0.08, 0.19 |  | 0.06 | -0.14, 0.26 |  | 0.02 | -0.18, 0.22 |
| human interest (looking to human) | 0.01 | -0.12, 0.13 |  | 0.00 | -0.19, 0.18 |  | 0.01 | -0.17, 0.20 |
| human interest (interacting with human) | 0.09 | -0.05, 0.22 |  | 0.08 | -0.12, 0.27 |  | -0.05 | -0.25, 0.15 |
| cylinder task (fast and accurate) | -0.07 | -0.21, 0.07 |  | -0.13 | -0.33, 0.07 |  | -0.13 | -0.32, 0.07 |
| cylinder detour (fast and accurate) | **0.18** | 0.06, 0.29 |  | 0.09 | -0.09, 0.28 |  | 0.18 | -0.01, 0.36 |
| unsolvable (looking to human) | 0.08 | -0.06, 0.22 |  | 0.01 | -0.20, 0.21 |  | -0.01 | -0.21, 0.19 |
| unsolvable (independent problem solving) | -0.11 | -0.24, 0.02 |  | 0.04 | -0.16, 0.24 |  | -0.01 | -0.20, 0.19 |

**Table S11.** Associations between dam C-BARQ scores at approximately 1 year of age and subsequent maternal behaviour. CI = Credible Interval.

|  | Week 1 | |  | Week 2 | |  | Week 3 | |
| --- | --- | --- | --- | --- | --- | --- | --- | --- |
| **Dependent Measure** | **Beta** | **90% CI** |  | **Beta** | **90% CI** |  | **Beta** | **90% CI** |
| attachment & attention-seeking | 0.01 | -0.17, 0.19 |  | -0.12 | -0.41, 0.17 |  | 0.06 | -0.21, 0.32 |
| chasing | -0.05 | -0.25, 0.14 |  | **-0.46** | -0.74, -0.17 |  | -0.17 | -0.45, 0.11 |
| dog-directed fear | 0.03 | -0.15, 0.21 |  | -0.12 | -0.41, 0.18 |  | 0.05 | -0.22, 0.31 |
| energy level | 0.13 | -0.04, 0.30 |  | -0.16 | -0.45, 0.13 |  | 0.04 | -0.23, 0.32 |
| excitability | **-0.17** | -0.34, -0.01 |  | **-0.28** | -0.55, -0.002 |  | -0.21 | -0.46, 0.04 |
| non-social fear | 0.05 | -0.11, 0.22 |  | 0.05 | -0.24, 0.33 |  | -0.03 | -0.28, 0.22 |
| separation-related behavior | -0.10 | -0.30, 0.10 |  | -0.30 | -0.62, 0.02 |  | -0.13 | -0.43, 0.15 |
| stranger-directed aggression | 0.06 | -0.11, 0.23 |  | -0.09 | -0.38, 0.21 |  | -0.06 | -0.31, 0.18 |
| stranger-directed fear | -0.05 | -0.23, 0.14 |  | -0.22 | -0.51, 0.07 |  | -0.17 | -0.43, 0.08 |
| touch sensitivity | -0.01 | -0.19, 0.16 |  | -0.05 | -0.34, 0.24 |  | -0.08 | -0.35, 0.18 |
| trainability | 0.13 | -0.06, 0.31 |  | **0.33** | 0.04, 0.61 |  | 0.09 | -0.18, 0.37 |

**Table S12**

Results from a linear mixed model predicting maternal behaviour principal component scores as a function of fixed effects for weeks of life, litter size, dam parity, breed composition, birth season, and rearing location.

| **Parameter** | **Beta** | **90% CI** |
| --- | --- | --- |
| week |  |  |
| 1 | — | — |
| 2 | -1.03 | -1.22, -0.85 |
| 3 | -1.55 | -1.73, -1.36 |
| litter size | -0.01 | -0.14, 0.12 |
| parity | -0.03 | -0.16, 0.10 |
| breed composition |  |  |
| 0-25% Labrador | — | — |
| 25-50% Labrador | 0.10 | -0.34, 0.53 |
| 50-75% Labrador | -0.27 | -0.84, 0.30 |
| 75-100% Labrador | -0.06 | -0.40, 0.27 |
| birth season |  |  |
| April To June | — | — |
| January To March | **0.39** | **0.05, 0.73** |
| July To September | 0.19 | -0.14, 0.54 |
| October To December | **0.47** | **0.14, 0.81** |
| location |  |  |
| Breeder Caretaker | — | — |
| Canine Early Development Center | 0.09 | -0.16, 0.33 |
| Abbreviations: CI = Credible Interval | | |

**Table S13**

Comparison to chance performance for DCDB object choice tasks among all puppy participants.

| **Variable** | **Null hypothesis** | **Mean** | **t** | **df** | **p** |
| --- | --- | --- | --- | --- | --- |
| communicative marker | 50 | 69.87 | 15.73 | 228 | 0.00 |
| arm pointing | 50 | 64.12 | 11.54 | 225 | 0.00 |
| odor control trials | 50 | 49.16 | -0.74 | 223 | 0.46 |
| memory (5s) | 50 | 75.33 | 19.35 | 226 | 0.00 |
| memory (10s) | 50 | 67.93 | 12.95 | 224 | 0.00 |
| memory (15s) | 50 | 65.75 | 9.28 | 145 | 0.00 |
| memory (20s) | 50 | 63.53 | 5.66 | 84 | 0.00 |
| auditory discrimination | 50 | 61.06 | 9.51 | 233 | 0.00 |
| odor discrimination: first choice | 50 | 56.82 | 5.42 | 231 | 0.00 |
| odor discrimination: final choice | 50 | 70.76 | 15.85 | 231 | 0.00 |
| visual discrimination | 50 | 91.88 | 53.08 | 233 | 0.00 |

**Table S14.** Associations between maternal behaviour and puppy behavioural and cognitive outcomes. CI = Credible Interval.

|  | Week 1 | |  | Week 2 | |  | Week 3 | |
| --- | --- | --- | --- | --- | --- | --- | --- | --- |
| **Dependent Measure** | **Beta** | **90% CI** |  | **Beta** | **90% CI** |  | **Beta** | **90% CI** |
| retrieval | -0.06 | -0.19, 0.07 |  | 0.08 | -0.04, 0.20 |  | -0.05 | -0.17, 0.07 |
| communicative marker | -0.10 | -0.24, 0.03 |  | **-0.13** | -0.26, 0.00 |  | 0.08 | -0.06, 0.21 |
| pointing gesture | 0.07 | -0.06, 0.20 |  | 0.05 | -0.09, 0.17 |  | -0.03 | -0.17, 0.12 |
| visual discrimination | 0.05 | -0.08, 0.17 |  | -0.01 | -0.14, 0.11 |  | 0.02 | -0.11, 0.14 |
| auditory discrimination | **-0.14** | -0.27, -0.01 |  | **-0.16** | -0.28, -0.04 |  | -0.04 | -0.17, 0.09 |
| novel object (bold) | -0.03 | -0.15, 0.09 |  | 0.06 | -0.05, 0.17 |  | 0.02 | -0.09, 0.12 |
| surprising events (bold) | -0.12 | -0.24, 0.02 |  | -0.03 | -0.15, 0.09 |  | -0.02 | -0.15, 0.11 |
| laterality index (right preference) | -0.12 | -0.24, 0.00 |  | **-0.16** | -0.27, -0.04 |  | 0.04 | -0.07, 0.16 |
| laterality strength | -0.02 | -0.14, 0.10 |  | -0.03 | -0.15, 0.09 |  | 0.04 | -0.08, 0.16 |
| warmups (trials to criterion) | 0.02 | -0.11, 0.16 |  | **-0.16** | -0.29, -0.03 |  | 0.00 | -0.13, 0.13 |
| human interest (looking to human) | -0.04 | -0.18, 0.10 |  | -0.03 | -0.17, 0.10 |  | -0.02 | -0.16, 0.12 |
| human interest (interacting with human) | 0.07 | -0.05, 0.20 |  | -0.09 | -0.21, 0.04 |  | -0.03 | -0.16, 0.10 |
| odor discrimination (accurate and persistent) | 0.05 | -0.09, 0.18 |  | **-0.13** | -0.25, -0.01 |  | -0.10 | -0.22, 0.03 |
| cylinder task (fast and accurate) | **0.17** | 0.04, 0.29 |  | -0.04 | -0.16, 0.08 |  | -0.04 | -0.17, 0.09 |
| cylinder detour (fast and accurate) | 0.10 | -0.03, 0.22 |  | -0.03 | -0.14, 0.09 |  | -0.03 | -0.15, 0.09 |
| memory (accurate) | -0.11 | -0.23, 0.01 |  | 0.05 | -0.07, 0.17 |  | 0.09 | -0.03, 0.21 |
| unsolvable (looking to human) | 0.05 | -0.09, 0.18 |  | 0.08 | -0.04, 0.22 |  | 0.03 | -0.09, 0.16 |
| unsolvable (independent problem solving) | 0.00 | -0.13, 0.14 |  | -0.06 | -0.19, 0.07 |  | -0.04 | -0.18, 0.11 |

| **Table S15.** Associations between maternal behaviour in weeks 1-3 and offspring 6-month and 12-month C-BARQ scores. CI = Credible interval. | | | | | | | | | | | | | | | | | | |
| --- | --- | --- | --- | --- | --- | --- | --- | --- | --- | --- | --- | --- | --- | --- | --- | --- | --- | --- |
|  | | Week 1 maternal investment | | | | |  | Week 2 maternal investment | | | | |  | Week 3 maternal investment | | | | |
|  | | *6-month CBARQ* | |  | *1-year CBARQ* | |  | *6-month CBARQ* | |  | *1-year CBARQ* | |  | *6-month CBARQ* | |  | *1-year CBARQ* | |
| **Measure** | | **Beta** | **90% CI** |  | **Beta** | **90% CI** |  | **Beta** | **90% CI** |  | **Beta** | **90% CI** |  | **Beta** | **90% CI** |  | **Beta** | **90% CI** |
| attachment & attention-seeking | 0.06 | | -0.08, 0.20 |  | 0.12 | -0.02, 0.25 |  | 0.04 | -0.09, 0.16 |  | 0.05 | -0.07, 0.17 |  | -0.02 | -0.15, 0.10 |  | 0.00 | -0.12, 0.13 |
| chasing | | 0.12 | -0.01, 0.26 |  | **0.14** | 0.00, 0.28 |  | 0.08 | -0.06, 0.21 |  | 0.10 | -0.03, 0.22 |  | -0.11 | -0.25, 0.02 |  | 0.02 | -0.10, 0.15 |
| dog rivalry | | 0.00 | -0.14, 0.13 |  | -0.07 | -0.21, 0.07 |  | 0.01 | -0.12, 0.14 |  | 0.05 | -0.07, 0.17 |  | -0.04 | -0.17, 0.08 |  | 0.09 | -0.04, 0.22 |
| dog-directed aggression | | 0.07 | -0.07, 0.21 |  | 0.00 | -0.12, 0.13 |  | 0.13 | -0.01, 0.27 |  | -0.02 | -0.13, 0.10 |  | 0.09 | -0.05, 0.22 |  | -0.01 | -0.13, 0.11 |
| dog-directed fear | | 0.10 | -0.04, 0.25 |  | -0.04 | -0.17, 0.09 |  | **0.14** | 0.01, 0.27 |  | 0.03 | -0.09, 0.15 |  | 0.04 | -0.09, 0.16 |  | -0.04 | -0.17, 0.08 |
| energy level | | -0.04 | -0.18, 0.09 |  | 0.06 | -0.07, 0.19 |  | -0.05 | -0.17, 0.07 |  | 0.02 | -0.11, 0.14 |  | -0.03 | -0.15, 0.09 |  | **-0.12** | -0.25, -0.01 |
| excitability | | 0.03 | -0.10, 0.15 |  | 0.00 | -0.14, 0.13 |  | 0.04 | -0.09, 0.16 |  | 0.06 | -0.06, 0.18 |  | -0.06 | -0.18, 0.06 |  | -0.05 | -0.17, 0.07 |
| non-social fear | | **0.14** | 0.00, 0.27 |  | 0.10 | -0.03, 0.22 |  | 0.07 | -0.06, 0.20 |  | 0.05 | -0.08, 0.17 |  | -0.05 | -0.18, 0.08 |  | -0.07 | -0.20, 0.06 |
| owner-directed aggression | | 0.04 | -0.09, 0.16 |  | -0.06 | -0.18, 0.07 |  | 0.07 | -0.04, 0.19 |  | 0.00 | -0.12, 0.12 |  | **0.14** | 0.03, 0.26 |  | 0.09 | -0.03, 0.21 |
| separation-related behavior | | -0.04 | -0.18, 0.09 |  | 0.11 | -0.03, 0.24 |  | **0.14** | 0.02, 0.26 |  | **0.13** | 0.00, 0.25 |  | -0.01 | -0.13, 0.11 |  | **0.12** | 0.00, 0.24 |
| stranger-directed aggression | | -0.09 | -0.22, 0.04 |  | 0.02 | -0.12, 0.15 |  | 0.12 | -0.01, 0.25 |  | -0.05 | -0.17, 0.08 |  | 0.03 | -0.10, 0.16 |  | -0.09 | -0.21, 0.04 |
| stranger-directed fear | | 0.01 | -0.12, 0.14 |  | -0.06 | -0.21, 0.07 |  | **-0.15** | -0.27, -0.03 |  | -0.06 | -0.19, 0.07 |  | -0.06 | -0.18, 0.06 |  | -0.11 | -0.24, 0.02 |
| touch sensitivity | | 0.05 | -0.08, 0.17 |  | -0.04 | -0.17, 0.08 |  | 0.09 | -0.02, 0.22 |  | 0.10 | -0.02, 0.21 |  | 0.03 | -0.08, 0.15 |  | 0.08 | -0.04, 0.19 |
| trainability | | -0.09 | -0.22, 0.05 |  | -0.03 | -0.16, 0.09 |  | 0.01 | -0.12, 0.14 |  | -0.05 | -0.17, 0.07 |  | -0.04 | -0.17, 0.09 |  | -0.10 | -0.22, 0.01 |

**Table S16.** Associations between maternal behaviour in weeks 1-3 and offspring DIAS and Dog-ADHD Rating Scale scores at 10 and 16 months of age. As a reminder, the four DIAS principal components were Behavioral Regulation (dogs who scored high on this component were impulsive, excitable, impatient), Aggression (dogs who scored high on this component acted aggressively when excited or frustrated), Response to Novelty (dogs who scored high on this component were hesitant of and disinterested in novel things and situations), and Responsiveness (dogs who scored high on this component were easy to train, interested in new things, and agreeable). The two averaged scores on the Dog-ADHD rating scale were Inattention (dogs who scored highly tended to lose interest quickly, get easily distracted, have difficulty concentrating and listening, and were slow to learn, especially complex tasks) and Activity-impulsivity (dogs who scored highly were active, fidgety, and always on the move, as well as lacking self-control). CI = Credible Interval.

|  | Week 1 maternal behaviour | | | | |  | Week 2 maternal behaviour | | | | | |  | | Week 3 maternal behaviour | | | | |
| --- | --- | --- | --- | --- | --- | --- | --- | --- | --- | --- | --- | --- | --- | --- | --- | --- | --- | --- | --- |
|  | *10-month* | |  | *16-month* | |  | *10-month* | |  | *16-month* | |  | | *10-month* | | |  | *16-month* | |
| **Dependent Measure** | **Beta** | **90% CI** |  | **Beta** | **90% CI** |  | **Beta** | **90% CI** |  | **Beta** | **90% CI** |  | | **Beta** | | **90% CI** |  | **Beta** | **90% CI** |
| *DIAS* |  |  |  |  |  |  |  |  |  |  |  |  | |  | |  |  |  |  |
| aggression | -0.03 | -0.16, 0.10 |  | -0.02 | -0.15, 0.11 |  | **0.14** | 0.02, 0.26 |  | 0.11 | -0.01, 0.23 |  | | -0.01 | | -0.13, 0.12 |  | 0.06 | -0.06, 0.18 |
| behavioral regulation | 0.07 | -0.08, 0.21 |  | 0.06 | -0.07, 0.20 |  | 0.07 | -0.06, 0.20 |  | **0.13** | 0.00, 0.25 |  | | 0.08 | | -0.05, 0.21 |  | 0.12 | -0.01, 0.24 |
| response to novelty | **-0.14** | -0.27, -0.01 | | **-0.17** | -0.30, -0.04 |  | -0.08 | -0.20, 0.04 |  | -0.08 | -0.20, 0.04 |  | | 0.04 | | -0.08, 0.17 |  | -0.11 | -0.24, 0.01 |
| responsiveness | 0.03 | -0.11, 0.16 |  | 0.04 | -0.09, 0.17 |  | 0.02 | -0.10, 0.15 |  | -0.10 | -0.23, 0.02 |  | | -0.04 | | -0.17, 0.08 |  | **-0.19** | -0.32, -0.07 |
| *Dog-ADHD RS* |  |  |  |  |  |  |  |  |  |  |  |  | |  | |  |  |  |  |
| activity & impulsivity | 0.08 | -0.06, 0.22 |  | 0.01 | -0.12, 0.13 |  | 0.09 | -0.04, 0.22 |  | 0.05 | -0.08, 0.17 |  | | 0.11 | | -0.02, 0.24 |  | 0.01 | -0.12, 0.14 |
| inattention | 0.10 | -0.03, 0.23 |  | -0.01 | -0.14, 0.13 |  | 0.08 | -0.05, 0.21 |  | 0.04 | -0.08, 0.17 |  | | **0.14** | | 0.01, 0.26 |  | 0.07 | -0.05, 0.19 |

**Figure S1**

Study litter sample size by litter size and whelping location.

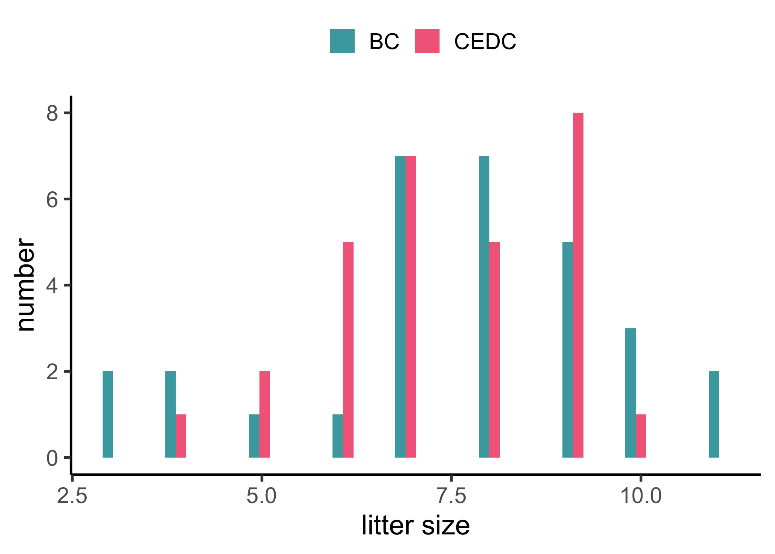

**Figure S2**

Principal component loadings for the two components of the cylinder task, inhibitory control trials and detour trials, derived from puppy performance. Positive component scores are associated with more accurate responses. For details on the underlying metrics and the experimental methods, see Gnanadesikan et al. (2023).

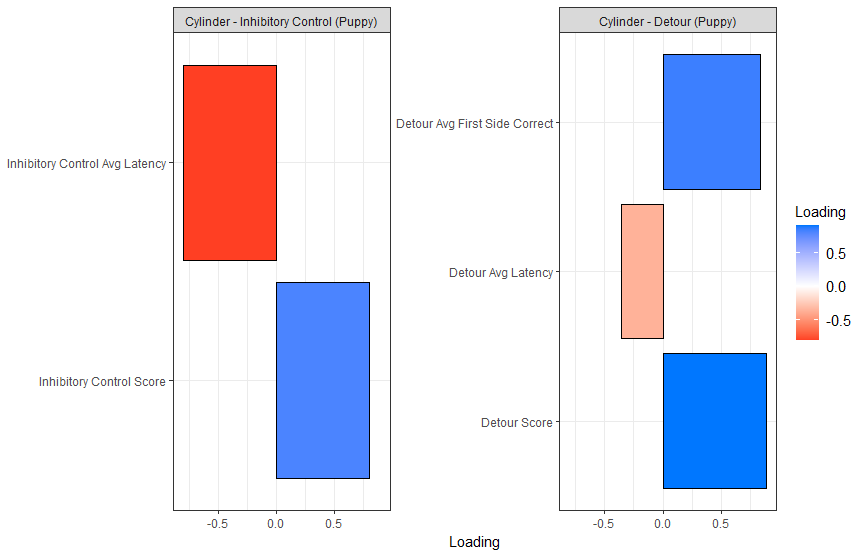

**Figure S3**

Principal component loadings for the two components of the cylinder task, inhibitory control trials and detour trials, derived from adult dog performance. Positive component scores are associated with more accurate responses. For details on the underlying metrics and the experimental methods, see Gnanadesikan et al. (2023).

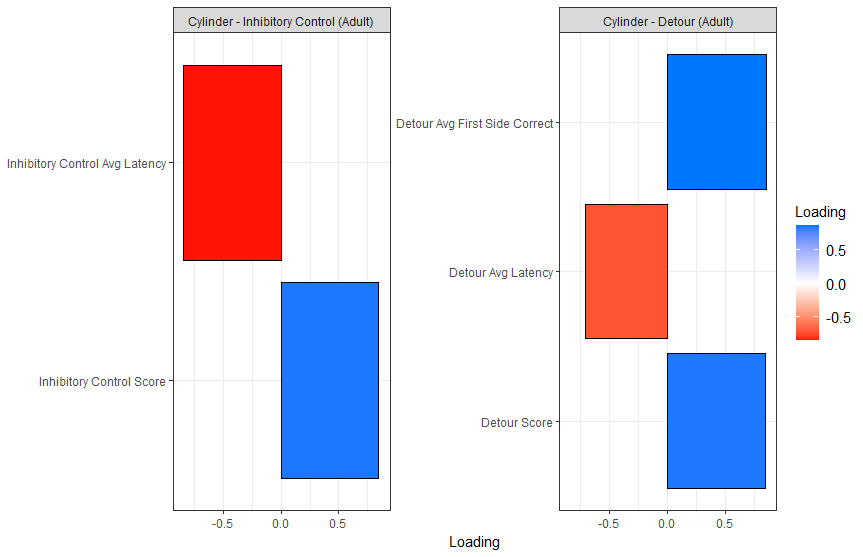

**Figure S4**

Principal component loadings for the odor discrimination task, derived from puppy performance. Positive component scores are associated with more accurate responses. For details on the underlying metrics and the experimental methods, see Gnanadesikan et al. (2023).

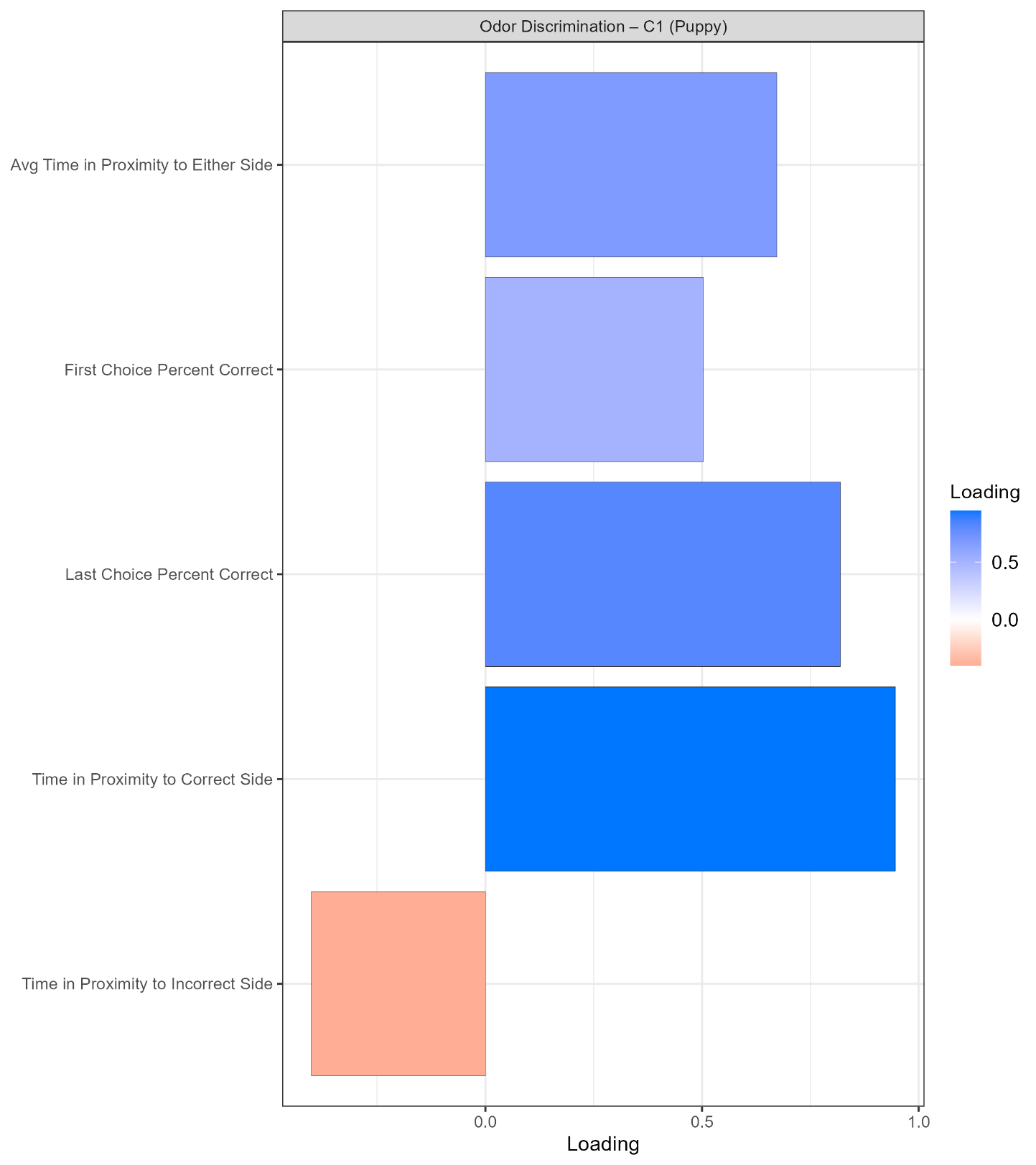

**Figure S5**

Principal component loadings for the novel object task, derived from puppy performance. Positive component scores are associated with more bold responses. For details on the underlying metrics and the experimental methods, see Gnanadesikan et al. (2023).

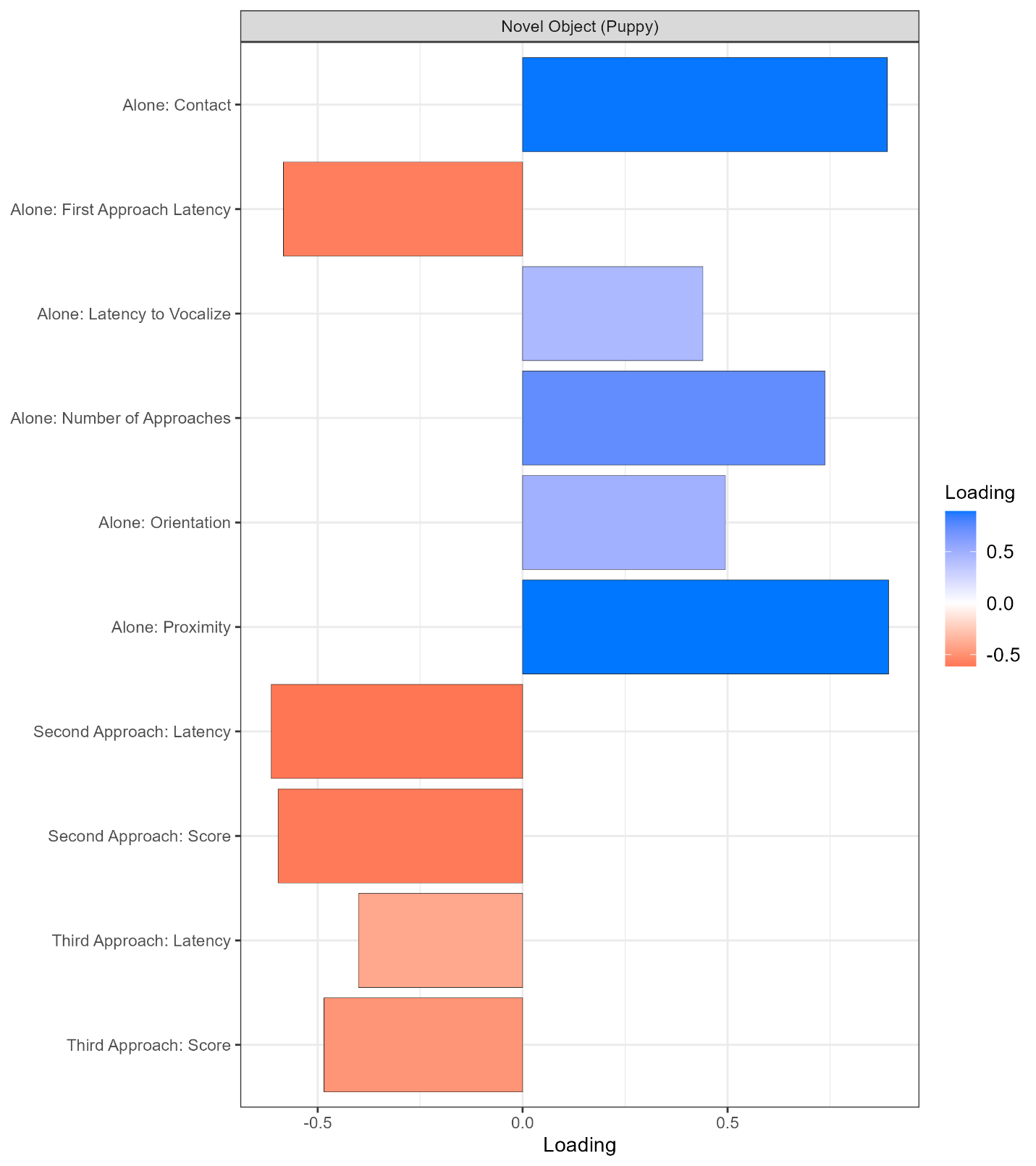

**Figure S6**

Principal component loadings for the novel object task, derived from adult performance. Positive component scores are associated with more bold responses. For details on the underlying metrics and the experimental methods, see Gnanadesikan et al. (2023).

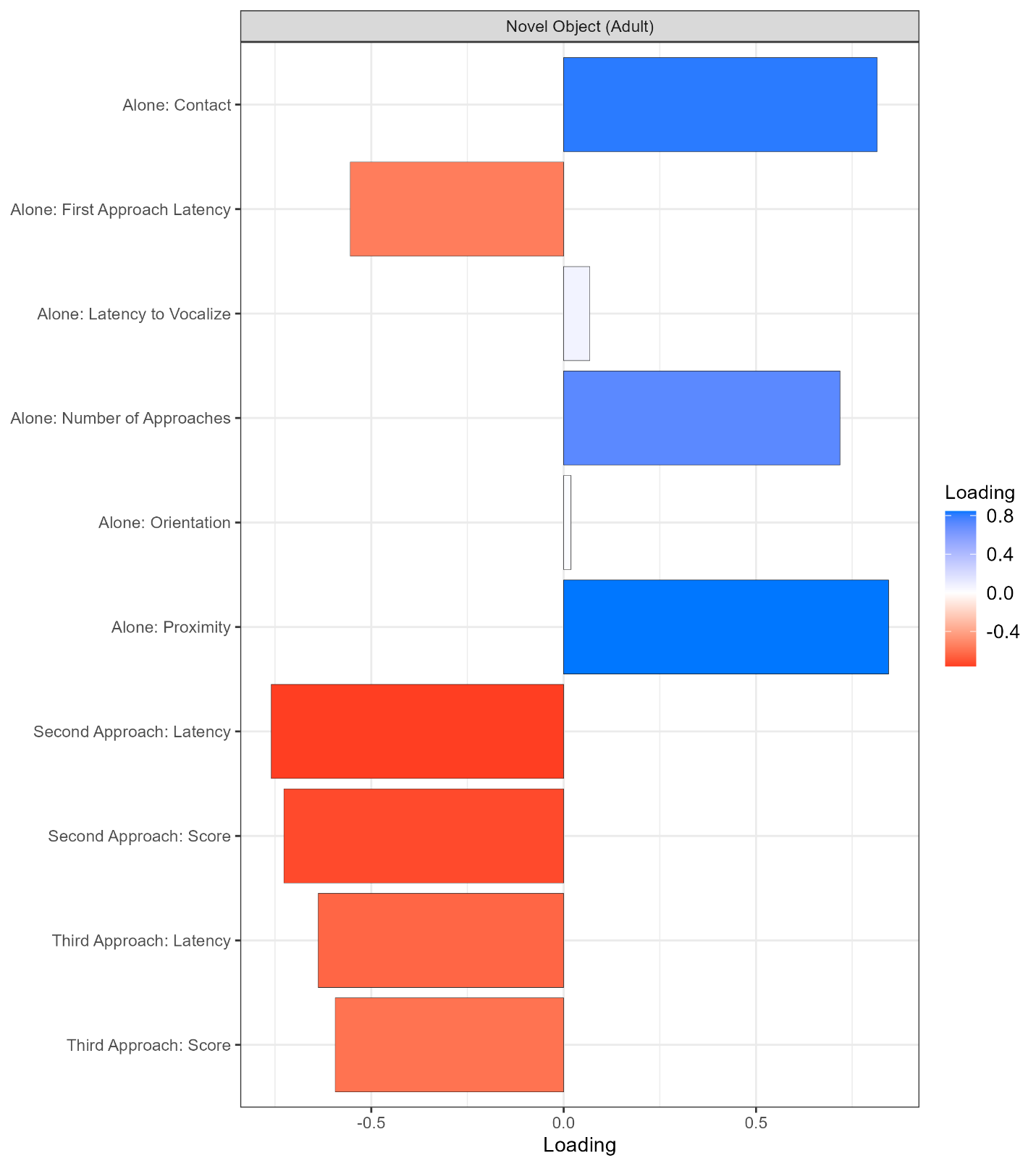

**Figure S7**

Principal component loadings for the surprising events task, derived from puppy performance. Positive component scores are associated with more bold responses. For details on the underlying metrics and the experimental methods, see Gnanadesikan et al. (2023).

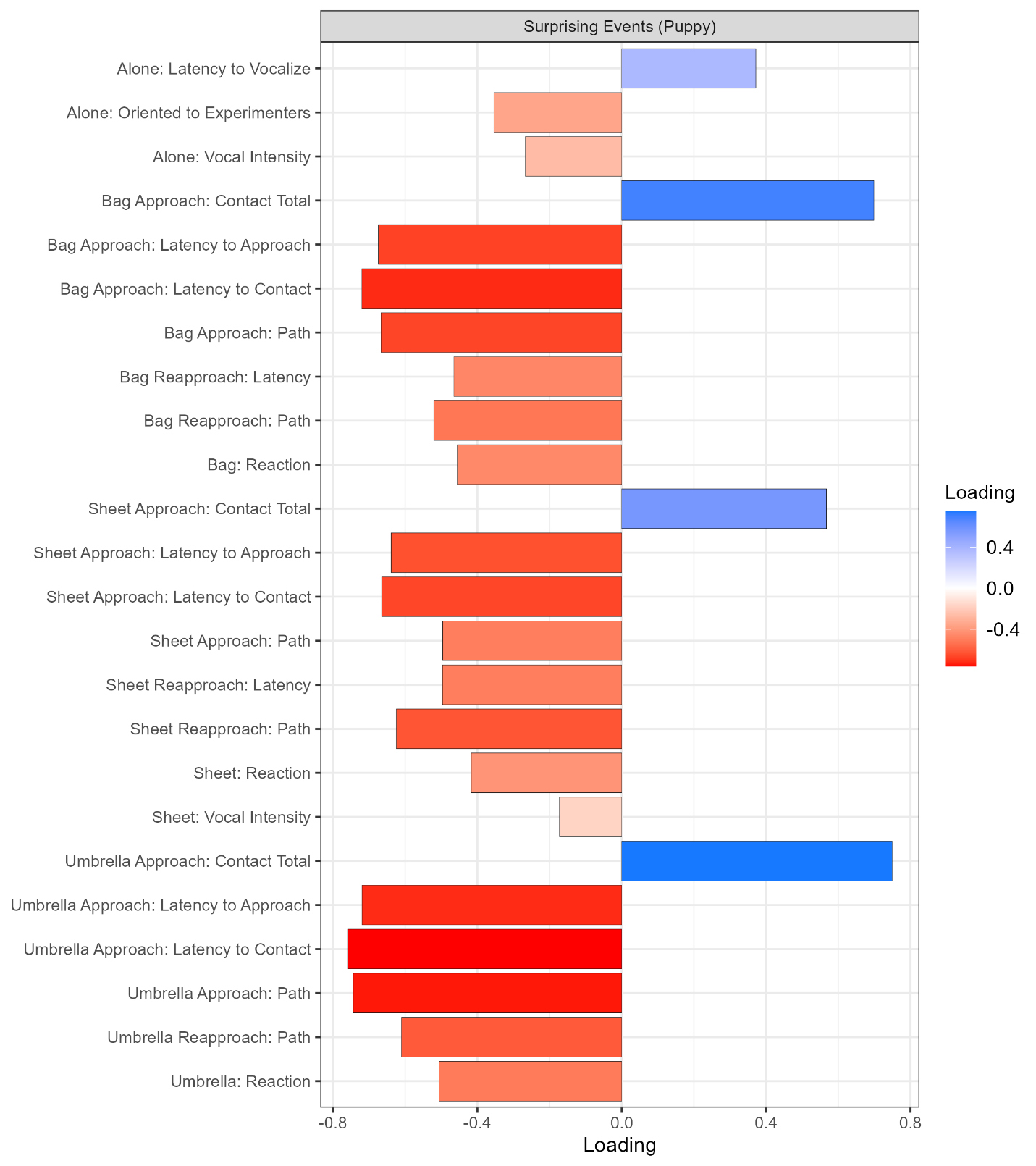

**Figure S8**

Principal component loadings for the surprising events task, derived from adult performance. Positive component scores are associated with more bold responses. For details on the underlying metrics and the experimental methods, see Gnanadesikan et al. (2023).

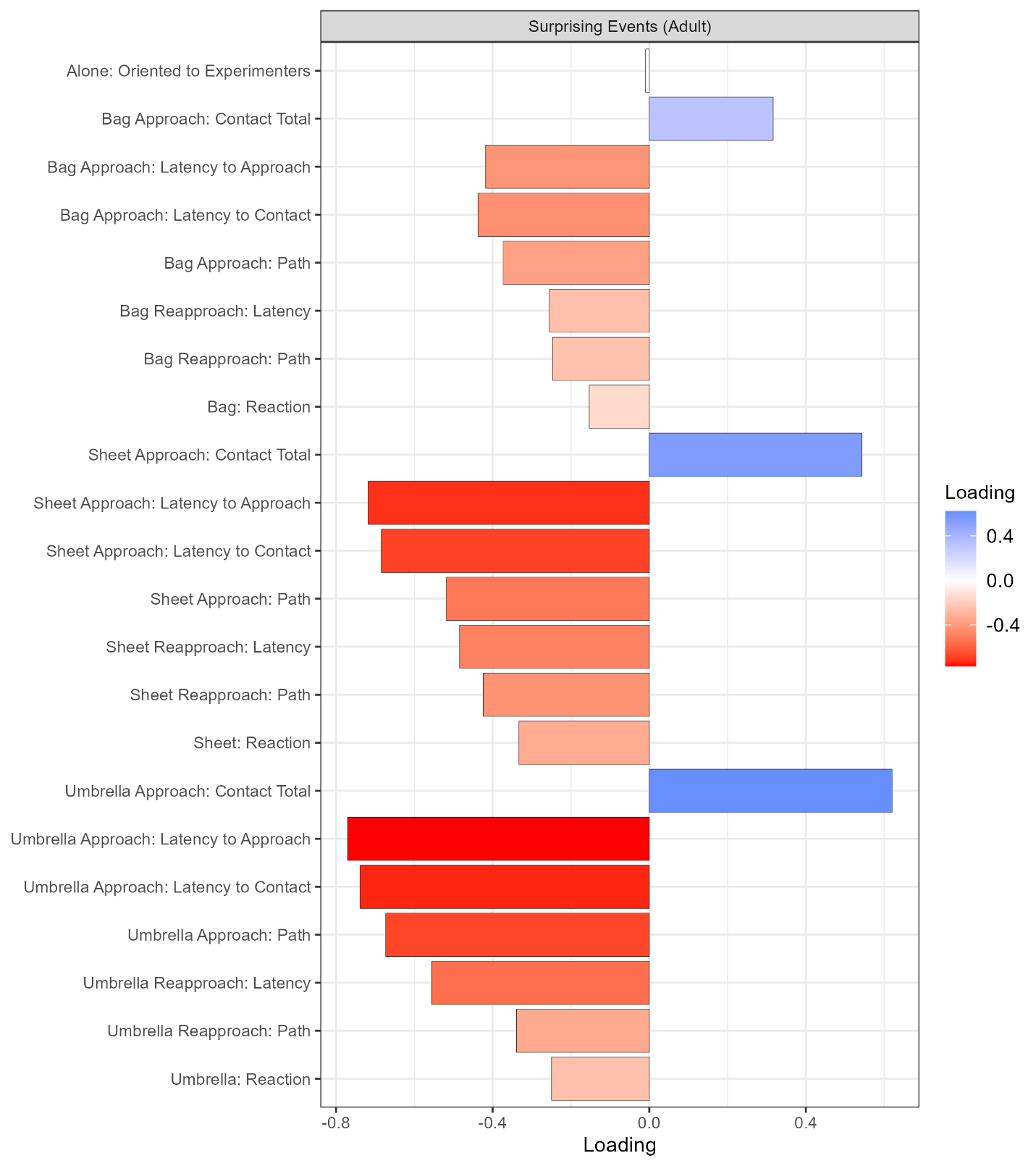

**Figure S9**

Correlation between individual-level maternal behaviour scores across three weeks of observation, calculated as principal components using 2 h versus 12 h of observations, in a subset of the sample. Sample size (*n*) and Pearson’s correlation coefficient (*r*) are reported in the panel.

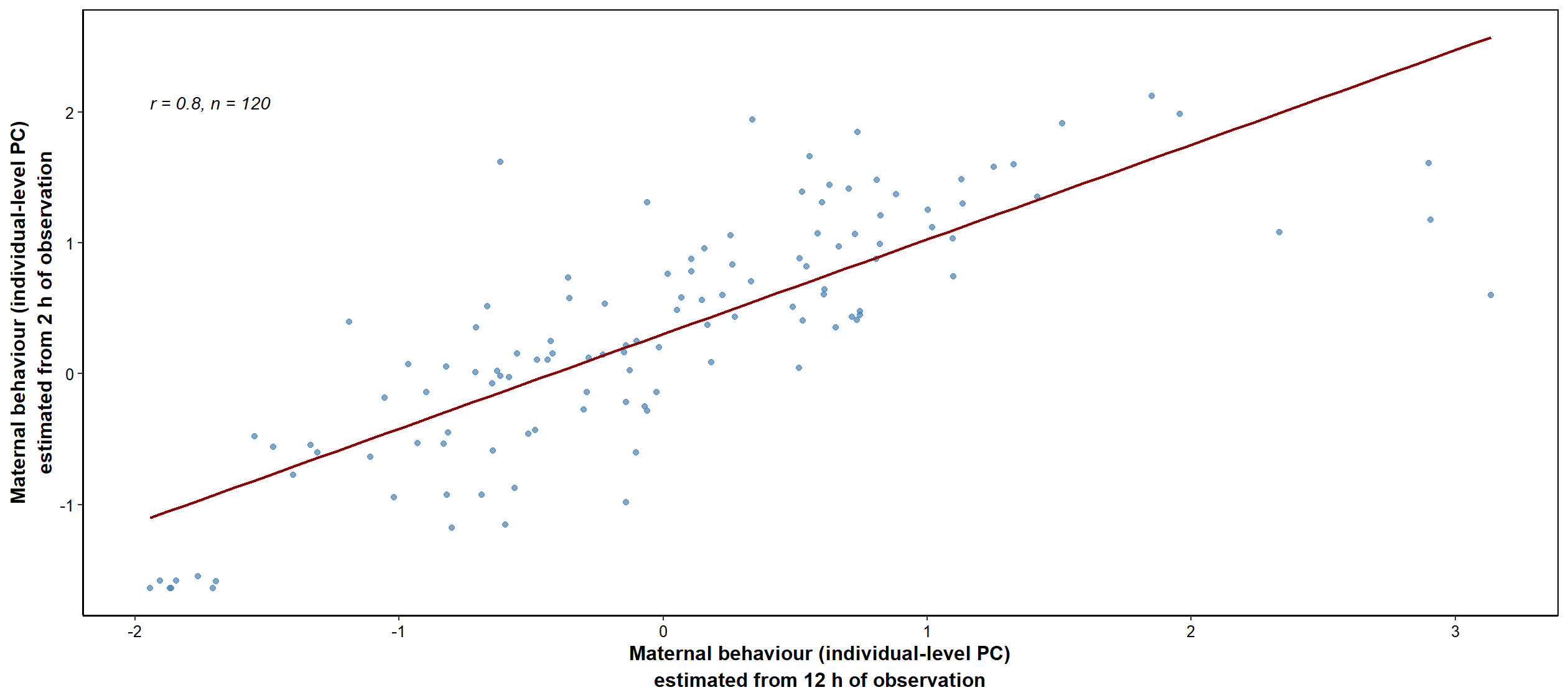

**Figure S10**

Correlation between litter-level maternal behaviour scores across three weeks of observation, calculated as principal components using 2 h versus 12 h of observations, in a subset of the sample. Sample size (*n*) and Pearson’s correlation coefficient (*r*) are reported in the panel.

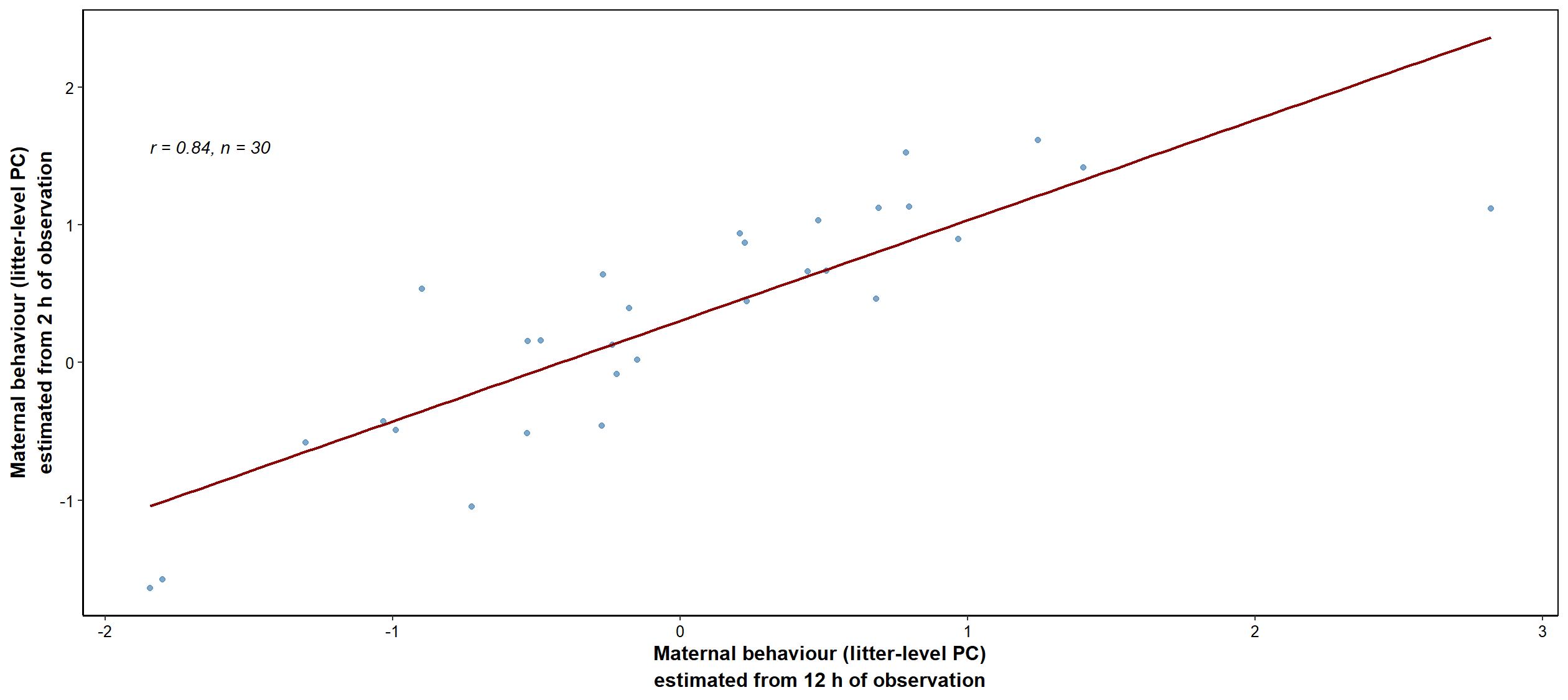
